# EHMT2 Controls Transcriptional Noise and the Developmental Switch after Gastrulation in the Mouse Embryo

**DOI:** 10.1101/2021.03.29.437567

**Authors:** Tie-Bo Zeng, Nicholas Pierce, Ji Liao, Purnima Singh, Wanding Zhou, Piroska E. Szabó

## Abstract

Embryos that carry zygotic or parental mutations in *Ehmt2*, the gene encoding the main euchromatic histone H3K9 methyltransferase, EHMT2, exhibit variable developmental delay. We asked the question whether the delayed embryo is different transcriptionally from the normally developing embryo when they reach the same developmental stage. We collected embryos carrying a series of genetic deficiencies in the *Ehmt2* gene and performed total RNA sequencing of somite stage-matched individual embryos. We applied novel four-way comparisons to detect differences between normal versus deficient embryos, and between 12-somite and 6-somite embryos. Importantly, we also identified developmental changes in transcription that only occur during the development of the normal embryo. We found that at the 6-somite stage, gastrulation-specific genes were not precisely turned off in the *Ehmt2*^−/−^ embryos, and genes involved in organ growth, connective tissue development, striated muscle development, muscle differentiation, and cartilage development were not precisely switched on in the *Ehmt2*^−/−^ embryos. Zygotic EHMT2 reduced transcriptional variation of developmental switch genes and at some repeat elements at the six-somite stage embryos. Maternal EHMT2-mutant embryos also displayed great transcriptional variation consistent with their variable survival, but transcription was normal in developmentally delayed parental haploinsufficient embryos, consistent with their good prospects. Global profiling of transposable elements in the embryo revealed that specific repeat classes responded to EHMT2. DNA methylation was specifically targeted by EHMT2 to LTR repeats, mostly ERVKs. Long noncoding transcripts initiated from those misregulated ‘driver’ repeats in *Ehmt2*^−/−^ embryos, and extended to several hundred kilobases, encompassing a multitude of additional, similarly misexpressed ‘passenger repeats.’ These findings establish EHMT2 as an important regulator of the transition between gastrulation programs and organ specification programs and of variability.

## INTRODUCTION

The development of the mammalian embryo entails chromatin remodeling and requires the activity of epigenetic modifiers. These effects are complex with regard to the timing and the source of the protein, for example whether it is passed on from the oocyte or produced in the embryo. Some of these epigenetic modifiers are known to play important roles at the early embryonic stages --in the oocyte, zygote, cleavage stages, and implantation-- as revealed by maternal mutation experiments that inactivate the genes in the oocyte (Fig. 1A, compare cross B versus control cross A) (1–14). Not only the absence, but also a reduced or increased level of these epigenetic modifiers in the female or male gamete may have lasting effects on the offspring (Fig. 1A, cross C versus cross A and cross D versus cross A). For example, paternal haploinsufficiency of *Setdb1*, encoding an H3K9 methyltransferase, correlates with a paternal effect of phenotype (15), revealing that a reduced dose of SETDB1 in the male gametogenesis affects gene expression in the offspring. Similarly, a paternal effect was found when overexpressing the H3K4 demethylase KDM1A during spermatogenesis resulting in reduced H3K4 methylation. This severely impaired development and survivability of the offspring and affected its transcriptome (16). Zygotic mutation experiments (Fig. 1A, cross E) gave evidence that chromatin regulators affect transcription and development at the postimplantation and fetal stages (17–19). It is not normally considered that the wild type offspring out of two heterozygous parents (Fig. 1A cross E) is biparental (maternal and paternal) haploinsufficient. We took advantage of mouse genetics to design a mouse study that allows measuring the effect of all these different deficiencies (Fig. 1A) on the development and the genome-wide transcription in the embryo. We included zygotic, maternal, and maternal-zygotic mutant embryos in one comprehensive experiment together with mono- or biparental haploinsufficient wild type individuals and wild type individuals from wild type parents (Fig. 1A). We focused on the deficiencies of the euchromatic histone H3 lysine-9-methyltransferase 2 (*Ehmt2*) gene.

**Figure 1.**
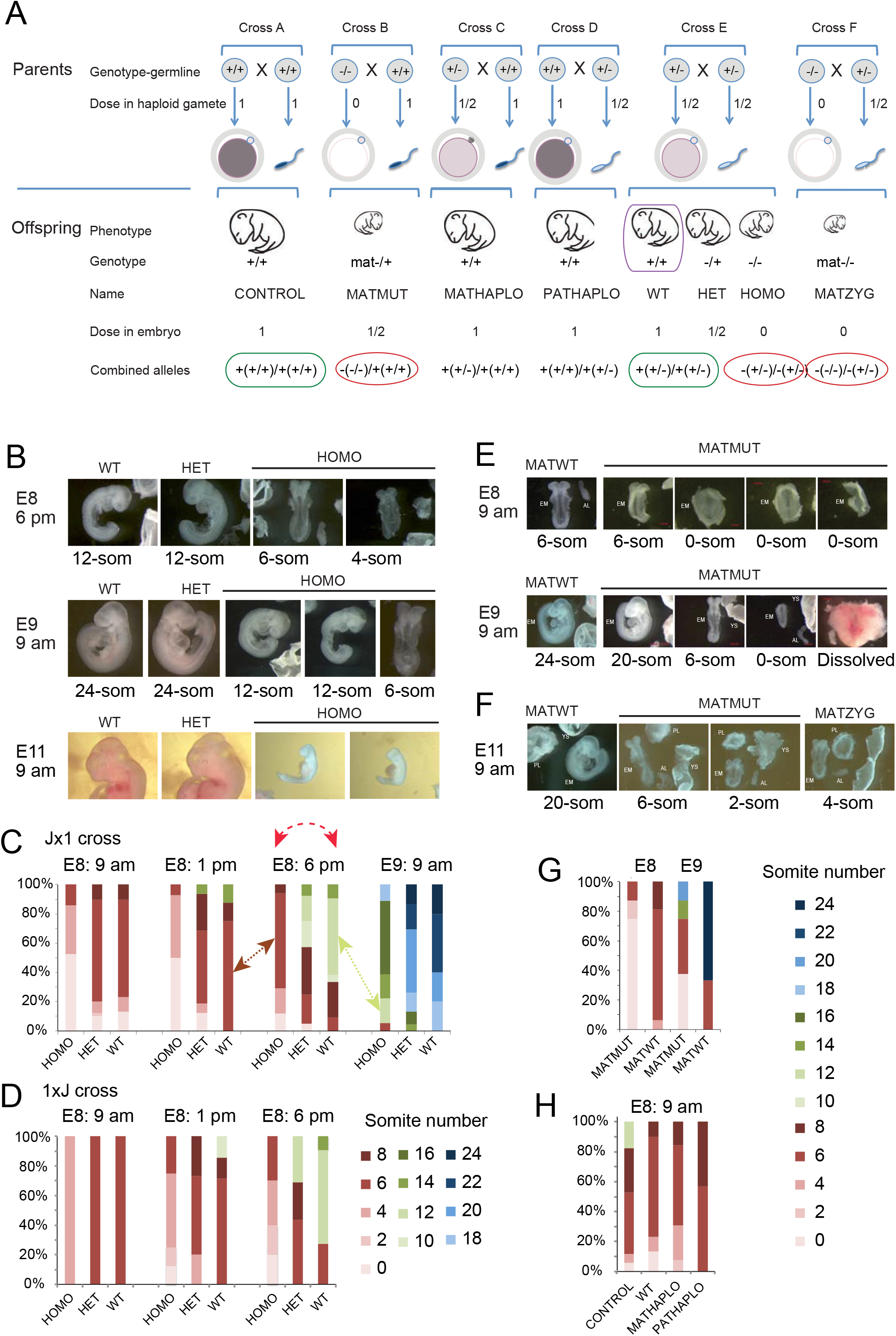
Zygotic and parental genetic deficiencies in *Ehmt2* result in delayed embryo development. (**A**) Study design. Genetic crosses to result in parental genotypes, and embryonic genotypes with insufficiencies of the epigenetic modifier in the gametes or in the embryo. We summarize the genetic crosses, the genotype of the parents (their diploid germ line and haploid gamete), the dose of the epigenetic factor in the gametes and in the embryo and the genotype of the embryo. We also provide the full allelic composition, which includes parental effects: allele from mother (mother’s genotype)/allele from the father (father’s genotype). Note that the WT embryo from the conventional test cross (purple box) is biparentally haploinsufficient. (**B-D**) Zygotic effect. (**B**) Images of representative embryos from crossing *Ehmt2*^+/−^ parents are shown, with the embryonic days and times of collection marked to the left. The embryo genotypes are marked at the top of each image and the somite numbers are marked underneath each image. (**C**) Percentages of the embryos that have reached specific somite numbers at collection are tallied from the *Ehmt2*^+/−^ x *Ehmt2*^+/−^ cross in the JF1×129 parental strain background. Day and time of collection is indicated above the bars. Red dashed arrow shows that comparing uterus-mate embryos according to conventional methodology means comparing more advanced wild type and delayed mutant embryos, and each group has its variation. The black arrow indicates comparing somite-matched embryos. The number of pregnant mothers and embryos were: E8: 9 am (n=17, 91), E8: 1 pm (n=7, 38), E8: 6 pm (n=14, 78) and E9: 9 am (n=7, 46). (**D**) Percentages of the embryos are similarly tallied in the reciprocal 129xJF1 parental strain background from the *Ehmt2*^+/−^ x *Ehmt2*^+/−^ cross. The number of pregnant mothers and embryos were: E8: 9 am (n=1, 8), E8: 1 pm (n=4, 30), and E8: 6 pm (n=5, 37). (D-F) Maternal effect. (**E**) Images of the *Ehmt2*^*m*−/z+^ embryos from the *Ehmt2*^fl/fl^; *Zp3-Cre^Tg/0^* x *Ehmt2*^+/+^ mating at E8: 9 am and E9: 9 am. (**F**) Representative images of MATMUT and MATZYG embryos at E9: 9 am. (n=5, 14). (**G**) Percentage of MATMUT and MATWT embryos at the somite stages as shown by colors (to the right). E8: 9 am (n=2, 7) and E9: 9 am (n=2, 6). (**H**) Parental haploinsufficiency and developmental delay. Tally of *Ehmt2*^+/+^ MATHAPLO embryos from the *Ehmt2*^+/−^ x *Ehmt2*^+/+^ cross (n=6, 25), and the *Ehmt2*^+/+^ PATHAPLO embryos from the *Ehmt2*^+/+^ x *Ehmt2*^+/−^ cross (n=4, 30), compared with CONTROL embryos from the *Ehmt2*^+/+^ x *Ehmt2*^+/+^ cross (n=5, 17) and WT embryos from the *Ehmt2*^+/−^ x *Ehmt2*^+/−^ cross (n=17, 91).

The *Ehmt2* gene encodes EHMT2, the major euchromatin-specific H3K9 histone dimethyltransferase (20), with a complex effect on embryonic development. *Ehmt2*^−/−^ zygotic mutant embryos die around 10.5 days post coitum (dpc) without exception (19, 21). Unlike the zygotic deficiency, the maternal mutation does not lead to fully penetrant lethality despite a strong maternal phenotype in the majority of the embryos, (2). It is not known whether *Ehmt2* paternal or maternal or biparental haploinsufficiencies have any long-lasting effect on embryo development or transcription. Important questions arise, such as why some maternal mutants survive but all zygotic mutants die; can the delayed development be correlated with changes in the transcriptome; is there a difference in developmental potential between a delayed and not delayed embryo in the absence of transcriptome differences.

To reveal the function of specific regulators, genetic experiments are employed to identify differentially expressed (DE) genes between wild type and mutant siblings. Such changes are generally interpreted to be the result of the mutation and are also considered to play a role in the phenotype. One caveat of such experiments is that if the mutation causes a delay in developmental progression, the differences between control and mutant uterus-mate embryos may largely result from comparing two embryos at different developmental stages. Delayed embryo development was reported in *Ehmt2*^−/−^ zygotic mutant embryos at 6.5 dpc (22) and at 7.5 dpc (23). *Ehmt2*^mat-/+^ maternal mutant embryos show partially penetrant developmental arrest at the 2-cell stage (2). *Ehmt2*^mat-/+^ maternal mutant embryos that pass the 2-cell stage exhibit a mild delay at the 8-cell stage, before destabilization of the inner cell mass lineage of the blastocyst occurs (14). We also noticed developmental delay in our *Ehmt2* knockout mouse line (24), which renders EHMT2 catalytically inactive, and is used in the current study.

Understanding developmental delay and its variability is an exciting question and requires a novel experimental approach. To investigate the role of EHMT2 during embryo development, we collected embryos carrying a series of *Ehmt2* deficiencies (Fig. 1A) between 8.5-9.5 dpc, shortly before dying, and performed total RNA sequencing of somite stage-matched individual embryos. We found differences not only in the transcriptome of normal versus deficient embryos, but also in the developmental changes in transcription that accompany normal versus deficient embryo development. We found the maternal and zygotic EHMT2 to be important in reducing transcriptional variation of genes and repeat elements at the six-somite stage embryos.

## RESULTS

We hypothesized that transcription changes shortly before death of the *Ehmt2* maternal and zygotic mutations will inform us what causes developmental delay or lethality in the embryos of different deficiencies in the genetic series (Fig. 1A). To characterize the developmental consequences of the deficiencies and to find the optimal collection times of somite-stage matched embryos, we first carried out time course experiments.

### Embryo developmental progression shows dose response to zygotic loss of EHMT2

We examined the *Ehmt2*^−/−^ homozygous (HOMO) embryos from the *Ehmt2*^+/−^ X *Ehmt2*^+/−^ parents (Fig. 1A cross E) at different time points and found that they exhibited delayed development compared to their *Ehmt2*^+/+^ wild type (WT) and *Ehmt2*^+/−^ heterozygous (HET) uterus mates (Fig. 1 B-D). Figure 1B displays representative images of uterus-mates with different genotypes. At 6 pm on embryonic day 8, the HOMO embryos only reached the 4 or 6 somite stage when the WT and HET embryos have reached the 12-somite stage. At 9 am on day 9, the HOMO embryos only reached the 12-somite stage while the WT and HET embryos have reached the 24-somite stage. We tallied the results of a larger number of pregnancies at 9 am, 1 pm and 6 pm on day 8, and at 9 am on day 9 of gestation as a percentage of each developmental stage in the total number of embryos collected per time point (Fig. 1C). We found that while the majority of WT embryos have reached the 6-somite stage by 9 am on day 8, HOMO embryos reached it at 6 pm on day 8. The majority of WT embryos have reached the 12-somite stage by 6 pm on day 8, whereas none of the HOMO embryos reached this somite number at that time. The embryos shown in Figure 1C were obtained by crossing *Ehmt2*^+/−^ mothers in the JF1/Ms genetic background and *Ehmt2*^+/−^ fathers in the 129S1 genetic background (Jx1 cross). A reciprocal cross (1×J) was also set up to validate the developmental phenotype, and to allow us to carry out allelespecific analysis of the transcriptome in the embryos based on SNPs between the mouse strains 129S1 and the JF1/Ms (Fig. 1D). We found that the developmental delay of HOMO embryos was reproducible in the reciprocal crosses (Fig. 1C-D). The tallies revealed an additional observation, which was not apparent at the time of dissections, that the development of HET embryos was slightly slower than the wild type embryos but faster than HOMO suggesting that developmental progression in the embryo was responsive to the dose of EHMT2.

### Embryo developmental progression shows dose response to maternal loss of EHMT2

To test the effect of the maternal *Ehmt2* mutation on development, we crossed *Ehmt2^fl/fl^; Zp3-Cre^Tg/0^* females with wild type males. The *Zp3-cre* transgene excises the floxed SET domain-coding exons of *Ehmt2* in the growing oocytes. The resulting maternal mutant *Ehmt2*^*mat*−/+^ (MATMUT) embryos lack maternal EHMT2 proteins in the oocyte, zygote, 1-cell, and early two-cell stages, but regain zygotic EHMT2 from the normal paternal allele, inherited from the sperm, after the onset of zygotic genome activation. Control *Ehmt2*^*fl*/+^ embryos (MATWT) were obtained from *Ehmt2^fl/fl^* mothers in the absence of cre. We found that the MATMUT embryos lagged behind the MATWT embryos, as illustrated in Figure 1E. The delay was even more drastic in maternal-zygotic *Ehmt2*^*mat*−/−^ (MATZYG) mutant embryos (Fig. 1F), which were obtained by crossing *Ehmt2^fl/fl^; Zp3-Cre^Tg/0^* females with *Ehmt2*^+/−^ males. The two most advanced MATZYG embryos we were able to recover had 6 somites (not shown) or 4 somites (Fig. 1F) at 9 am on day 11 of gestation.

We tallied the dissected embryos at 8.5 and 9.5 dpc (Fig. 1G). Whereas the mode of somite number was 6 for the MATWT embryos at 9 am on day 8, the MATMUT embryos only reached the 6-somite stage one day later, at 9 am on day 9, when the mode was 24 for the MATWT embryos. We also noticed that the developmental delay of MATMUT embryos varied greatly. At 9 am on day 9 some of these were at the presomite stage but some others displayed 20 somites. Despite the developmental delay observed in the somite-stage embryos, some MATMUT embryos developed to term, reached adulthood, and even reproduced successfully. We obtained live pups by crossing *Ehmt2^fl/fl^*; *Zp3-Cre*^*Tg*+^ females with males of three different wild type strains and with *Ehmt2*^+/−^. The average fecundity rate was 1.7 pups per plug (from 24 plugs total) as compared to 8.5 pups per plug in the control females (Table S1). This is in agreement with another study that obtained 2.8 MATMUT pups compared to 8.5 control pups (2). It is noteworthy that the gestation time of the MATMUT embryos was 20.5 days instead of the normal 19.5 days. MATMUT female adults gave birth to an average of 7 pups (from 5 plugs total). This result suggests that the developmental roadblock of MATMUT embryos can be completely overcome in a stochastic manner.

### Embryo developmental progression shows dose response to maternal and paternal haploinsufficiency of EHMT2

Next we compared the development of *Ehmt2*^+/+^ wild type (WT) embryos derived from two *Ehmt2*^+/−^ heterozygous parents to the *Ehmt2*^+/+^ (CONTROL) embryos derived from *Ehmt2*^+/+^ parents (Fig. 1A cross E). The mode was 6 somites in both crosses at 9 am on day 8. While none of the WT embryos has reached the 12-somite stage, 20% of the CONTROL embryos did (Fig. 1H). Less of the former reached the 8-somite stage also. Because all of these embryos had two normal alleles of the *Ehmt2* gene, the developmental difference may be explained by genetic difference in the parents’ genotype, the WT embryos being biparental haploinsufficiencient. To further evaluate this possibility, we set up crosses to generate maternal or paternal haploinsufficient (MATHAPLO or PATHAPLO) *Ehmt2*^+/+^ embryos from one *Ehmt2*^+/+^ parent and one *Ehmt2*^+/−^ parent, the mother or the father, respectively (Fig. 1A crosses C and D). We tallied the developmental stage of the embryos (Fig. 1H) and found that, unlike some CONTROL embryos, none of the MATHAPLO, or PATHAPLO embryos reached the 12-somite stage at 9 am on day 8. This slight delay in both MATHAPLO and PATHAPLO cases, suggests that both the mother’s, and the father’s mutant genotype affected the development of the wild type embryo. This implies that the dose of the EHMT2 protein in the oocyte and the sperm plays a role in instructing developmental programs in the embryo.

### RNA-seq experiment to uncover the role of EHMT2 in advancing embryo development

From the combined results in Figure 1B-H we can see that the maternal, zygotic and parental haploinsufficient mutations of *Ehmt2* affect developmental progression differently. We hypothesized that the different *Ehmt2* deficiencies (Fig. 1A) display unique patterns in their transcriptomes, which would inform us about the specific roles of maternal and zygotic EHMT2 on embryo development, and collected 6-somite and 12-somite stage embryos with the different embryo genotypes, and parental genotypes as listed in Table S2. The collection times were adjusted to the time course of the mutations. The HOMO embryos, for example, reached the mode of 6-somite stage by 6 pm on day 8, whereas the MATMUT embryos reached it at 9 am at day 9. The 1×J cross was slightly slower to develop than the Jx1 cross (Fig. 1C-D). To define the significance of EHMT2 in regulating developmental decisions we also collected WT *and* HOMO embryos at the 12-somite stage, which we will refer to as WT12 and HOMO12 to distinguish them from the 6-somite embryos, WT6 and HOMO6. We prepared total RNA from four replicates of individual embryos, two females and two males, in each condition, except for the very rare maternal-zygotic mutant embryos, which were duplicate females. We carried out allelespecific and strand-specific RNA sequencing analysis on total RNA samples as we did earlier using MEFs (25). We analyzed the results using three-way and four-way comparisons based on somite-matched embryos, revealing important false positives and false negatives of the conventional comparison that uses siblings.

### Three-way comparison differentiates true hits and false positives

A conventional comparison between uterus-mate HOMO and WT embryos will involve comparing HOMO6 and WT12 embryos at 8.5 dpc according to the modes of somite number at that time (dotted arrow in Fig. 1C). However, such WT12-HOMO6 comparison will result in a mixed outcome and will include two major components as depicted in Figure S1A: 1) “what it takes to be normal” at the 6-somite stage”, which can be assessed by comparing the WT6 versus HOMO6 samples and 2) “what it takes to develop” from the 6- to 12-somite stage, which can be assessed by comparing the WT12 versus WT6 samples. To be able to separate those components, we first performed a three-way comparison of uterus-mate HOMO6 and WT12 embryos, including WT6 embryos in the analysis (Fig. S1A). The HOMO6 and WT6 embryos had to be obtained from different dams and at different time points (black arrows in Figure 1C), but they were genetically identical due to the same inbred mouse lines used.

We identified differentially expressed (DE) genes between the three conditions. Each vector (colored arrows in Fig. S1A) was assigned a direction toward the healthier and/or more developed state, asking what is different in the WT versus HOMO embryos, and what is different in the 12-somite versus 6-somite stage.

Venn diagrams (Fig. S1B-C) depict the summary of the 3-way comparison (Table S3). We detected 388 genes that had decreased expression in WT12 versus HOMO6 embryos. The majority of these hits (225 out of 388) also decreased in the WT6 versus HOMO6 embryos (thick orange arrow in Fig. S1B), revealing that those transcripts are suppressed by EHMT2, and belong to the “what it takes to be normal” category. However, an additional 145 DEGs were decreased in the WT6-HOMO6 comparison, and these would be missed in the conventional WT12-HOMO6 comparison. Out of the 388 hits, 77 were found also in the WT12-WT6 comparison (open green arrow in Fig. S1B), reflecting developmental changes in transcription that occur between the 6- and 12-somite stages in normal embryos. These changes do not occur in response to the *Ehmt2* mutation, but are false positive hits of the WT12-HOMO6 comparison.

We detected 206 genes that showed increased expression in the WT12 versus HOMO6 embryos (Fig. S1C). This is surprising, considering that EHMT2 is a suppressor of gene activity. While only 24 out of 206 hits were detected in the intersection with the WT6-HOMO6 comparison (orange arrow in Fig. S1C), the majority (138 out of 206) occurred in the intersection with the WT12-WT6 comparison (open green arrow in Fig. S1C), revealing that those changes occur between the 6- and 12-somite stages during wild type embryo development, independent of EHMT2. These are false positive hits of the WT12-HOMO6 comparison. The false positive hits in Fig. S1B and S1C are further illustrated by examples, *Lhx1* and *Ascl1* (Fig. S1D-E), and heatmaps (Fig. S1F-G).

### EHMT2 is required for DNA CpG methylation at specific genes in the embryo

In addition to methylating H3K9, EHMT2 also affects transcription by regulating DNA methylation in the embryo, in male germ cells, and in ES cells at specific genes and transposable elements (22, 26–29). To find out how EHMT2 affects DNA methylation we performed whole genome bisulfite sequencing (WGBS) analysis of HOMO and WT embryo DNA at 9.5 dpc. We then identified differentially methylated regions (DMR) between the WT and HOMO samples, at DEGs, which have DNA methylation in either condition at the TSS (+/−1000 bp) (Table S4). The DMRs most frequently occurred at the intersection of WT12-HOMO6 and WT6-HOMO6 comparisons (Fig. S2A and S2B). We plotted the DMR DNA methylation and RNA expression levels of the DEGs from these intersections. DMRs occurred at 34 of 225 DEGs downregulated in the WT samples (Fig. S2C), such as *Naa11, Asz1, Pdha2*, and *Magea10, Hormad2*. In each case a higher level of DNA methylation corresponded to less RNA expression in the WT versus HOMO embryo, suggesting that EHMT2-dependent suppression at these genes involves DNA methylation in the WT embryos. On the other hand, 9 DMRs at 24 DEGs showed the opposite (Fig. S2D). A lower level of DNA methylation corresponded to higher RNA expression in the WT versus HOMO embryos, suggesting that EHMT2 helps to maintain the DNA hypomethylated state of these 24 genes in the WT embryo, most likely indirectly. Most of these genes were germ-line-specific genes. Examples of the above two classes are displayed by IGV browser images of *Hormad2* and *Zfp873* (Fig. S2E). Importantly, the WGBS analysis combined with the threeway comparison of RNA-seq results allowed us to conclude that EHMT2–dependent DNA methylation was more frequent at the DEGs in the “what it takes to be normal” than in the “what it takes to develop” category (50 versus 9 DEGs, respectively).

### Four-way comparison stratifies developmental changes in the *Ehmt2* mutant embryos

To further improve the interpretation of our RNAseq experiment we included one more condition, the 12-somite stage *Ehmt2*^−/−^ embryo (HOMO12) in our analysis. This four-way comparison (Fig. 2A) has two advantages over the three-way comparison: we can now also determine “what it takes to be normal” in the embryo at the 12-somite stage by contrasting WT12-HOMO12 and “what it takes to develop” normally or abnormally between the 6- and 12 somite stages by using the contrasts of WT12-WT6 and HOMO12-HOMO6 embryos.

**Figure 2.**
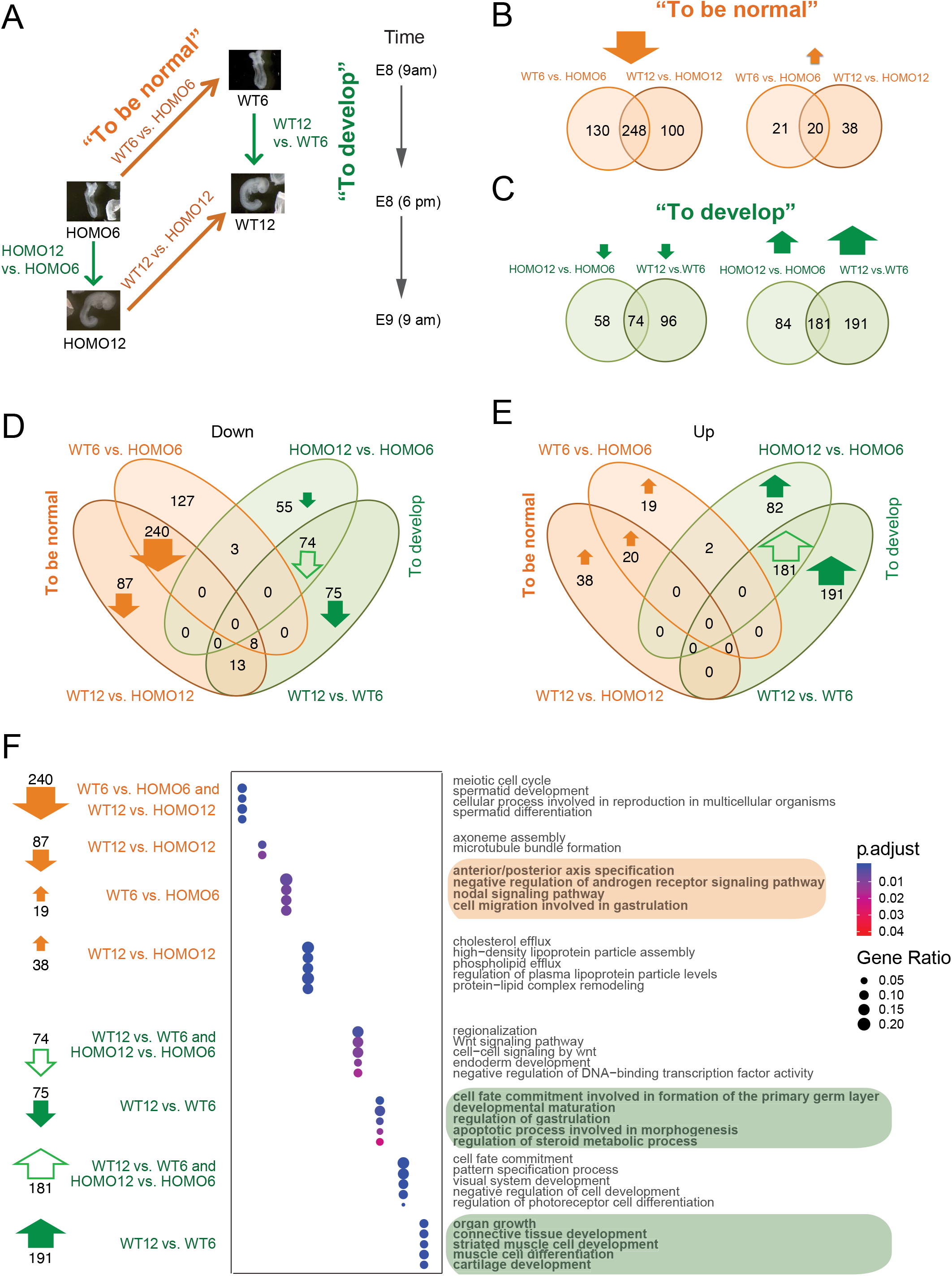
A four-way comparison identifies DEGs that distinguish normal embryo development from mutant embryo development. (**A**) Four-way comparison. The four questions asked are depicted by arrows: “what it takes to develop” from 6- to 12-somite stage in the normal or in the mutant embryo (green arrows indicate the WT12-WT6 and HOMO12-HOMO6 comparisons), and “what it takes to be normal” at 6-somites or at the 12-somite stage (orange arrows mark the WT6-HOMO6 and WT12-HOMO12 comparisons). (**B**) Identifying what it takes to be normal. Venn diagrams show the number of the downregulated (left) and upregulated (right) DEGs between the *Ehmt2*^+/+^ (WT) and the *Ehmt2*^−/−^ (HOMO) embryos at the 6-somite stage and at the 12-somite stage. (**C**) Identifying what it takes to develop from 6- to 12-somite stage. Venn diagrams show the number of downregulated (left) and upregulated (right) DEGs between the 12-somite and 6-somite stages, during normal *Ehmt2*^+/+^ (WT), or mutant *Ehmt2*^−/−^ (HOMO) development. (**D-E**) Venn diagrams merge all of the downregulated (**D**), or all of the upregulated (**E**) DEGs from the two-way comparisons as described above. (**F**) Gene ontology analysis of the DEGs revealed by the four-way comparison. Bubble plots depict the GO terms enriched in the comparisons as indicated at the top. Arrows highlight the DEGs from specific segments of the Venn diagrams.

To find out what it takes to be normal, we identified 378 downregulated DEGs in the WT embryos using comparisons of the WT6-HOMO6 and 348 between WT12-HOMO12 (Fig. 2B). 248 of these DEGs overlapped between the two contrasts. Compared to 478 downregulated genes, much fewer, only 79 DEGs were upregulated in the WT embryos in the totality of WT-HOMO comparisons. To find out what it takes to develop, we identified 372 and 265 upregulated DEGs at the 12-somite stage in the WT12-WT6 and HOMO12-HOMO6 contrasts, respectively. 181 DEGs sorted into the intersection (Fig. 2C). Compared to 456 upregulated genes, only 228 were downregulated in the totality of the 12-somite versus 6-somite comparisons. We found that “what it takes to be normal” are mainly DEGs that require EHMT2 for suppression (Fig. 2B), and “what it takes to develop” are DEGs that require EHMT2 for their activation along development (Fig. 2C).

To uniquely stratify these changes, we merged the Venn diagrams into downregulated (Fig. 2D), or upregulated (Fig. 2E) DEGs from the two-way comparisons as described above (Fig. 2B and 2C). It was interesting to note that, with very few exceptions, the DEGs populated the 6 compartments on the outside of the composite Venns, revealing that there is very little overlap between the DEGs that belong to “what it takes to be normal” at the left side and “what it takes to develop” to the right of each Venn. The diagrams illustrate also that the majority of DEGs are either downregulated in WT versus HOMO embryos (Fig. 2D) or upregulated in 12-somite versus 6-somite stage embryos (Fig. 2E). To understand what these changes mean, we performed a pathway analysis (Fig. 2F, and Table S5) of the DEGs (Table S3) from the prominent sections of the Venn diagrams. We also generated heatmaps and browser examples from these transcript sets (Fig. 3). First, we looked at the “what it takes to be normal” categories. The intersections of WT6-HOMO6 and WT12-HOMO12 contain 240 downregulated DEGs (Fig. 2D and Fig. 3A and B) that require EHMT2 for their suppression in WT embryos regardless of somite number of 6 or 12. These DEGs are enriched in gene ontology (GO) terms, such as meiotic cell cycle, spermatid development and differentiation, and cellular process involved in reproduction (Fig. 2F), including *Asz1, Dnmt3l, Sycp2, Rnf212, Syce1, Piwil4, Hormad2, Smcb1, Spaca1, Cftr, Zpbp, Ccdc42*, and the *Xlr3-Xlr5* cluster (*Xlr4c, Xlr3a, Xlr3c, Xlr3b, Xlr4a, Xlr4b, Xlr5a, Xlr5c*, and *Xlr5b*) (Table S3). The 20 genes upregulated in both the WT6-HOMO6 and WT12-HOMO12 comparisons (Fig. 2E and Fig. 3C-D) include DEGs, such as *Sycp1, Tssk6, Il3ra*, and *Csf2ra* (Table S3) and genes GO terms, such as sperm chromatin condensation, spermatid nucleus differentiation, reciprocal meiotic recombination, homologous recombination, and synapsis (Fig. 2F and Table S5).

**Figure 3.**
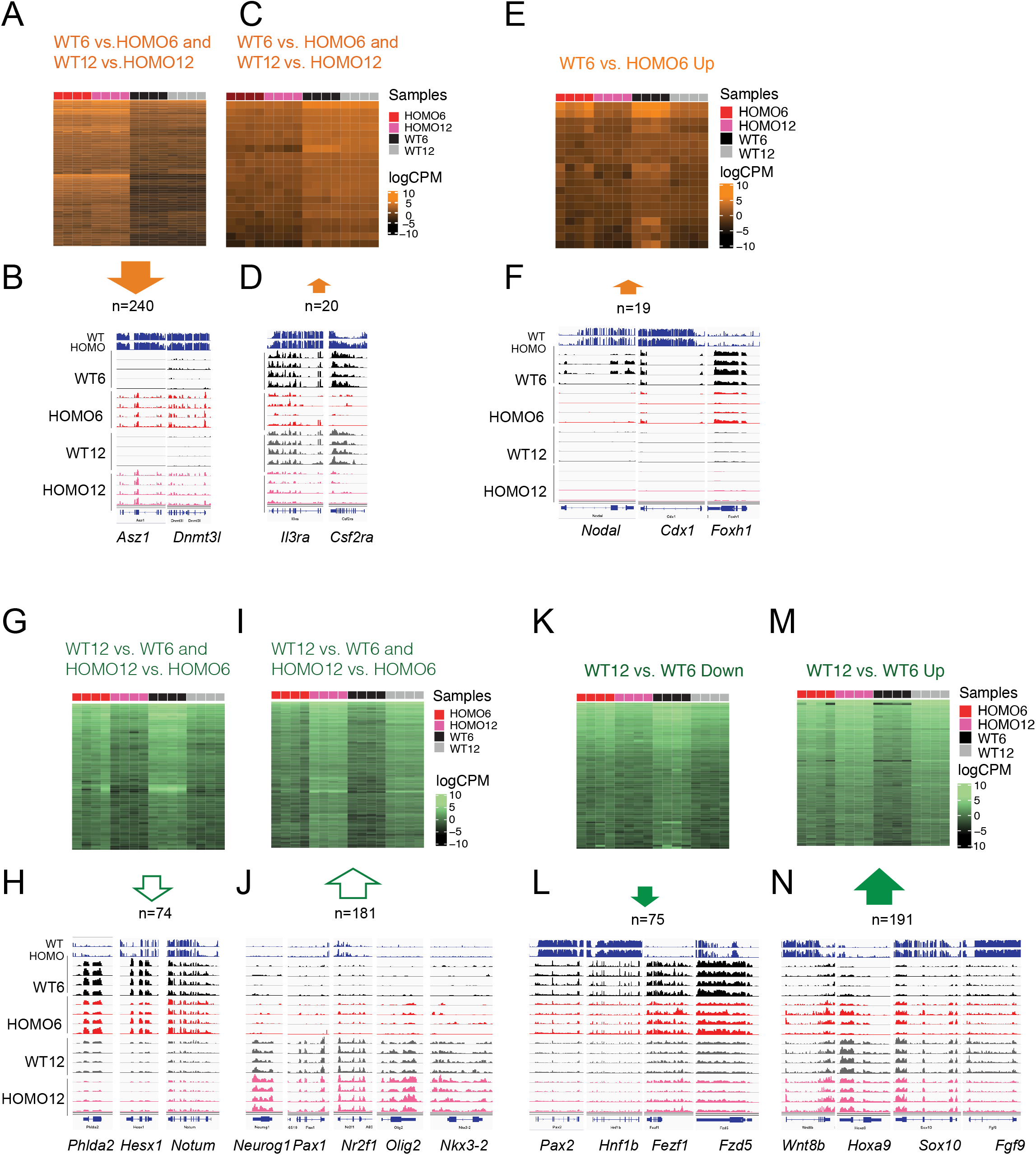
What it takes to be normal and what it takes to develop. (**A-F**) **What it takes to be normal.** Changes occur in response to EHMT2 at both the 6-somite and 12-somite stages (**A-D**) Changes detected in response to EHMT2 between WT6 vs. HOMO6, but not between WT12 vs. HOMO12 embryos (**E-F**). (**G-N**) **What it takes to develop.** (**G-J**) Developmental DEGs are downregulated (A) or upregulated (C) between the 6-and 12-somite stages regardless of EHMT2. (**K-N**) Downregulation of 75 genes (E) or upregulation of 191 genes (G) only occurs only in normal development (WT12 vs. WT6), but not in mutant development (HOMO12 vs. HOMO6) between the 6-and 12-somite stages; therefore, these developmental-specific changes require EHMT2. These are false negatives of the conventional WT12-HOMO6 comparison. Heatmaps (**A, C, E, G, I, K, and M**) are shown for the highlighted Venn-diagram sections as written and marked with the thick arrows. IGV browser examples (**B, D, F, H, J, L and N**) are shown for the DEGs that drive the GO term in the highlighted Venn-diagram sections.

The WT12-HOMO12 comparison includes 87 downregulated genes (Fig. 2D), such as *Dnah1a*, and *Dnah1b* and GO terms, such as axoneme assembly and microtubule bundle formation (Fig. 2F). The WT12-HOMO12 comparison includes 38 upregulated genes (Fig. 2E), such as *Apoa1, Apoa4, Apoc2* and *Apom* and terms related to protein-lipid complex remodeling (Fig. 2F). The WT6-HOMO6 downregulated genes had no significant GO terms. We found 19 upregulated genes in the WT6-HOMO6 comparison, such as *Lefty1, Nodal, Cdx1*, and *Foxh1* (Fig. 2E and 3E-F), which belong to developmental processes, such as anterior-posterior axis specification, negative regulation of androgen signaling pathway, nodal signaling, and cell migration involved in gastrulation (Fig. 2F, Table S5). Their expression drops between the 12-somite to 6-somite stages, and this occurs slower in WT6 versus HOMO6 embryos, suggesting that EHMT2 affects this process.

Next, we inspected the pathways associated with DEGs in the “what it takes to develop” categories. The intersections WT12-WT6 and HOMO12-HOMO6 comparisons inform us what it takes to develop from 6-to 12-somite stage in the WT and also in the mutant embryos. We found 74 DEGs in the intersection that are suppressed in the 12-somite stage compared to the 6-somite conditions (Fig. 2D and Fig. 3G-H), such as *Pou5f1, Cfc1, Zic3, Ar, Eomes, Gsc, Foxh1, Wnt8a, Cdx1, Amer3, Notum, Lect2, Sfrp5, Phlda2, Hesx1*, and *Pgr*. The GO terms, such as regionalization, Wnt signaling pathway, and endoderm development were enriched (Fig. 2F). We found 181 DEGs in the intersection that are upregulated in the 12-somite stage compared to the 6-somite conditions (Fig. 2E, and Fig. 3I-J), such as *Hoxa10, Hoxa9, Hoxa10, Hoxd10, Hoxd11, Lhx6, Dlx1, Dlx2, Neurod1, Neurod4, Neurog1, Myl2, Cyp26b1, Dbx1, Nr2f1, Nr2f2, Nr2e1, Sostdc1, Gdf7, Gdnf, Nkx2-1, Nkx3-2, Olig2, Sox8, Meox2, Pax1, Alx4, Uncx, Onecut1*, and *Ascl1*. These DEGs were associated with GO terms, such as pattern specification, cell fate commitment, and visual system development (Fig. 2F and Table S5). The intersections of WT12-WT6 and HOMO12-HOMO6 reflect developmental changes that occur in WT and HOMO embryos equally, undisturbed by the mutation. The HOMO12-HOMO6 comparison revealed 55 downregulated and 82 upregulated DEGs (Fig. 2D and E) not in an intersection of the Venn diagram. These DEGs did not result in any significantly enriched GO term.

The DEGs found in the WT12-WT6, but not in an intersection of the Venn diagram (Fig. 2D and E) are very interesting, because these reveal changes that occur consistently in normal development but not in the *Ehmt2*^−/−^ mutant embryo development between the 6- and 12 somite stages. The 75 DEGs downregulated in WT12-WT6 but not HOMO12-HOMO6 contrast (Fig. 2D) include *Nanog, Nodal, Mesp1, Pax2, Fzd5, Hnf1b*, and *Fezf1* and the GO terms such as cell fate commitment involved in formation of the primary germ layer, developmental maturation, regulation of gastrulation, and apoptotic process involved in gastrulation (Fig. 2F and Table S5). The 191 DEGs upregulated in WT12-WT6 but not in HOMO12-HOMO6 contrast (Fig. 2E) include *Erbb4, Neurog2, Wnt7b, Wnt8b, Myf5*, and *Fgf9*, and the GO terms such as organ growth, connective tissue development, striated muscle development, muscle differentiation, and cartilage development (Fig. 2F and Table S5). These developmental changes require EHMT2.

### EHMT2 suppresses variation of DEGs at the 6-somite stage

Looking at the WT12-WT6 segments of the Venn diagrams more closely in their heatmap dispalys, we noticed that even though we could only call the changes significant in the WT12-WT6 but not in HOMO12-HOMO6 comparisons, downregulation (Fig. 3K-L) or upregulation (Fig. 3M-N) of the DEGs did take place from the 6-to 12-somite stages in both WT and HOMO embryos. The heatmaps and browser images also showed that the HOMO6 embryos displayed the greatest variation among the samples (Fig. 3K-N). We concluded that this variation prevented us from calling statistically significant change in the HOMO12-HOMO6 comparison.

To explore EHMT2-dependent variation in more detail, we performed a principal component analysis (PCA) of the upregulated 191 DEGs and downregulated 75 genes (Fig. 4A-B) in the WT12-WT6 comparison, as defined in the Venn diagrams (Fig. 2D-E). Whereas WT6, WT12, HOMO12 samples are tightly clustered in the first and second principal components, the HOMO6 samples are scattered in both directions. In contrast, we did not find such scattering of the HOMO6 samples (Fig. 4C) in the PCA plot of the downregulated 240 DEGs from the intersection of WT6-HOMO6 and WT12-HOMO12 comparisons (Fig. 2D). This suggests, that EHMT2 more strongly affects the variation of DEGs, which “takes to develop normally” than those, which “takes to be normal”. We calculated the variance in each section of the 4-way Venn diagrams (from Fig. 2D-E) and found that the HOMO6 samples exhibit the greatest variability in 9 out of 12 comparisons (Fig. 4D). We also performed a mock comparison by substituting the HOMO6 1×J samples for the HOMO6 Jx1 samples and found that the HOMO6 samples were again the most variable in 9 out of 12 segments of the Venn diagram (Fig. 4D). In addition, Using MDSeq (30), we identified a set of genes, which are significantly (FDR < 0.05) differentially variable between the WT12 and HOMO6 embryos. The heatmap (Fig. 4E) shows the Z-scores of those genes in the HOMO6, HOMO12, WT6 and WT12 embryos. The HOMO6 embryos displayed the most heterogeneity. We showed earlier that the HOMO6 samples were the most variable among DEGs of the WT12-WT6 comparisons (Fig. 3K-N). In addition, the HOMO6 samples were variable in the group of developmental genes, such as *Nodal, Lefty1*, and *Foxh1*, which were downregulated from the 6- to the 12-somite stage in WT embryos but also belonged to the WT6-HOMO6 upregulated category. We found that, even though these genes exhibited tight regulation in WT6 embryos in each cross, they showed variability in the HOMO6 Jx1 and 1×J embryos (Fig. 4F), albeit at different levels of expression. In summary, EHMT2 suppresses the variation of developmental gene expression at the 6-somite stage.

**Figure 4.**
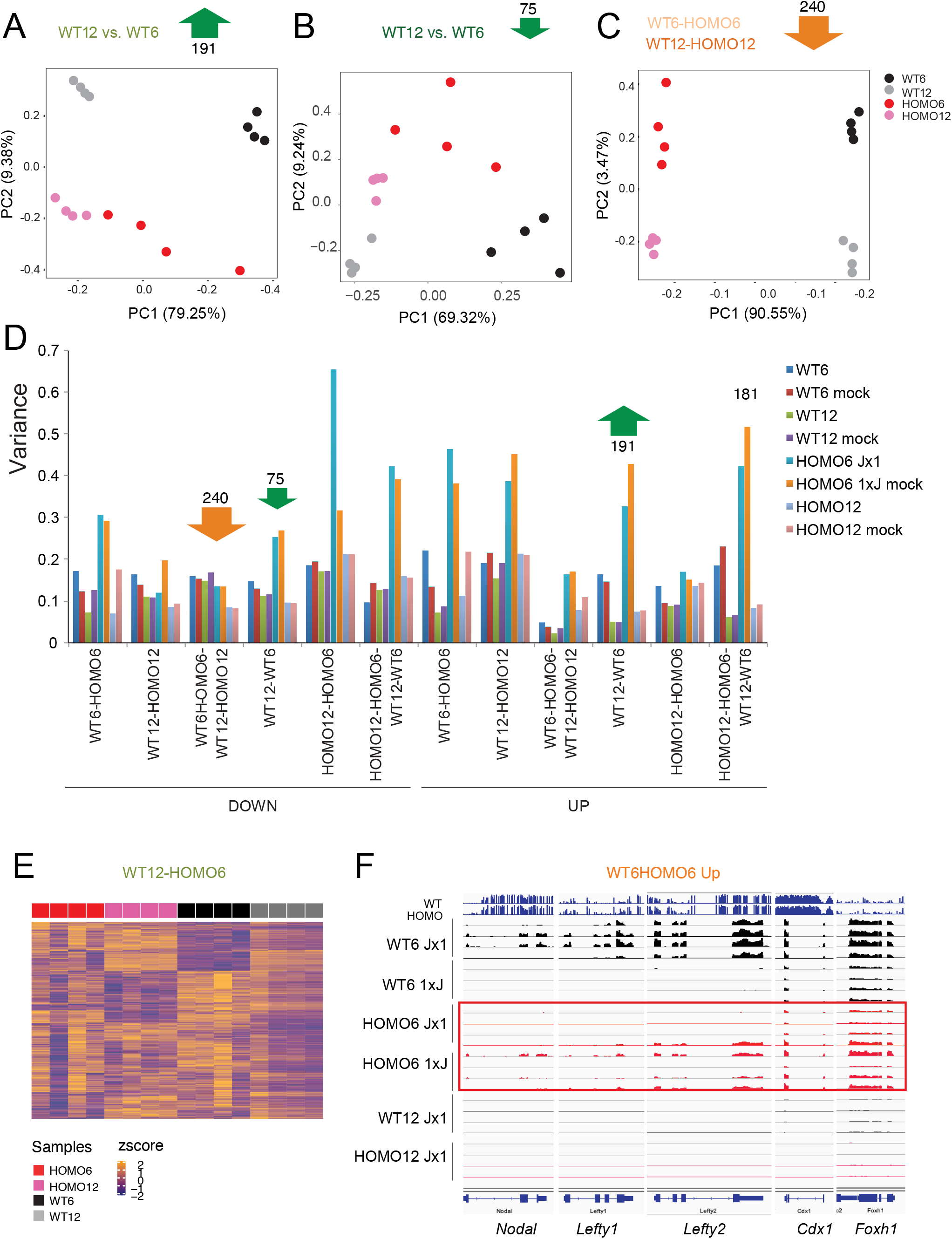
*Ehmt2* mutant embryos display an increased transcriptional variation. (**A-C**) Principal component analysis is shown for the samples marked to the right based on the DEGs derived from the four-way comparison as indicated at the top. (**D**) Bar graph displays the variance in the samples (to the right) for the DEGs in each comparison. The HOMO6 and WT6 samples from the reciprocal (1×J) cross were used to substitute for the (JX1 cross) in the four-way comparison to generate the mock results. The HOMO6 samples are the most variable. (**E**) Heatmap shows the z-scores of the genes that display the highest level of dispersion in the WT12-HOMO6 comparison based on MDSeq analysis. The HOMO6 samples are the most dispersed. (**F**) Variation is observed in a group of DEGs that belong to the WT6-HOMO6 comparison. IGV browser images of embryo samples are shown, four replicates in each condition as marked. The top two lanes are WGBS sequencing results.

### Expanding the findings to the entire set of genetic crosses

We carried out DEG calling (Table S3) in the entire dataset (Table S2) using the relevant comparisons. The MATHAPLO, PATHAPLO and WT (biparental haploinsufficient) samples only differed from the CONTROL samples, at a total 8, 7, and 4 DEGs (Fig. S3). 190 DEGs were called between the MATMUT6 embryos and MATWT6 embryos. We generated heatmaps (Fig. 5A and B) that show the transcriptional profile of each transcript out of the 6 prominent compartments of the Venn diagrams (as in Figure 2D and E) across each sample of our study. The categories of “what it takes to be normal” are seen at the top and the categories “what it takes to develop” are seen at the bottom. The transcriptome of MATZYG mutant embryos was very similar to the HOMO embryos in the WT6-HOMO6 and WT12-HOMO12 contrasts while it resembled the 6-somite embryo in the WT12-WT6 and HOMO12-HOMO6 contrasts. Overall, the zygotic mutation dominated the transcription pattern over the maternal mutation in these rare MATZYG embryos. The MATHAPLO, PATHAPLO and WT samples were very similar to the CONTROL samples. The MATMUT6 embryos displayed a highly variable pattern and resembled 12-somite embryos in the developmental comparisons.

**Figure 5.**
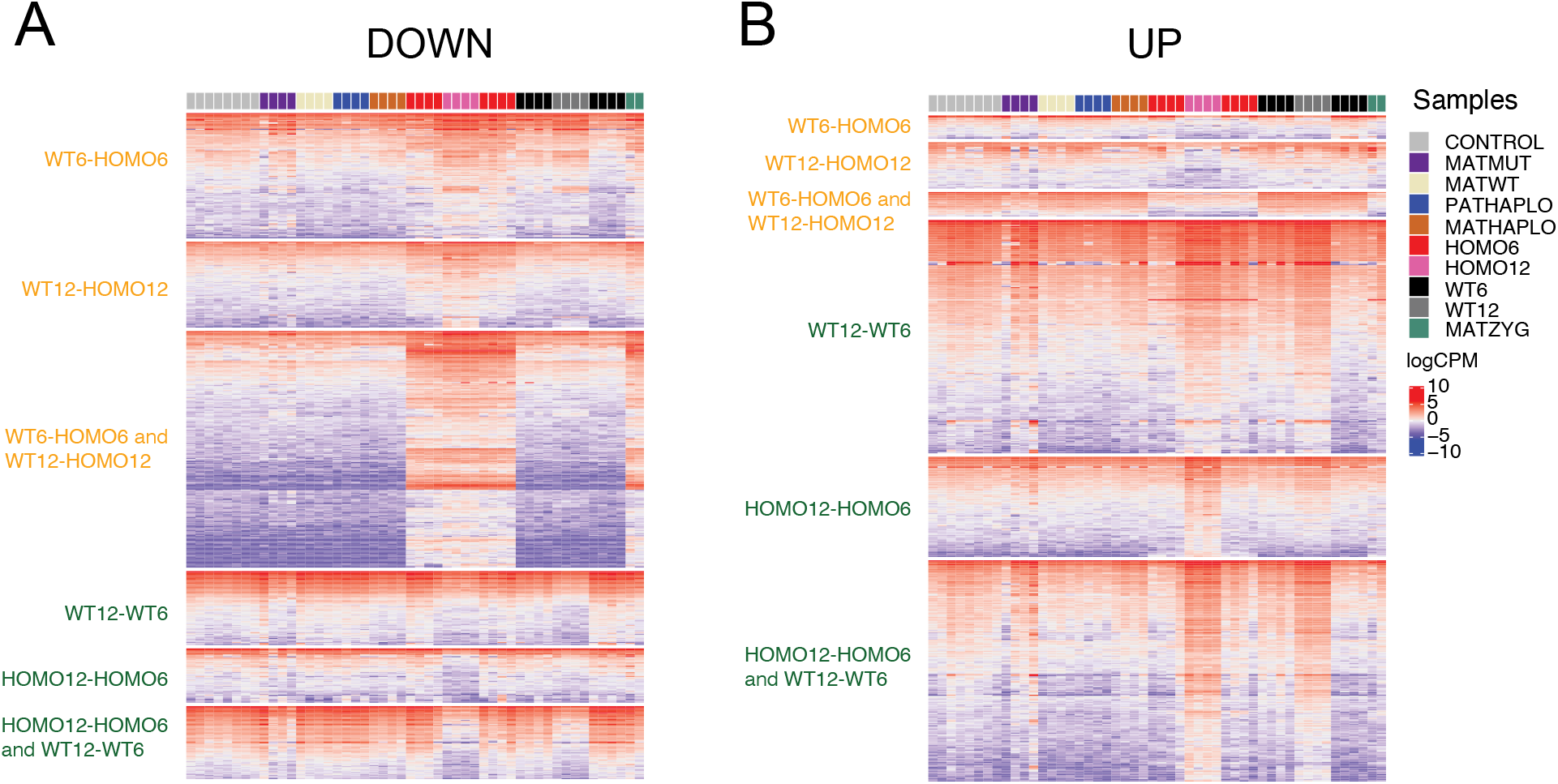
Transcriptomes of embryos with different genetic deficiencies. Heatmaps display gene expression values (logCPM) for the downregulated (**A**) and upregulated (**B**) genes from the compartments of the Venn diagrams in Figure 2 D-E, as marked, representing the full dataset (Fig. 1A and Table S2). Four replicates are shown in the samples coded by color (to the right).

### Maternal and paternal haploinsufficient embryos exhibit normal transcriptomes

To shed light on how the maternal and paternal haploinsufficiencies affect the transcriptome, we analyzed the RNAseq results of 6-somite stage MATHAPLO and PATHAPLO embryos. We found 8 DE transcripts in MATHAPLO embryos and 7 DEGs in PATHAPLO embryos compared to control 6-somite embryos (Table S3) and these did not provide GO enrichment. When we compared the biparental haploinsufficient (WT6) embryos to the corresponding CONTROL embryos derived from WT parents (Fig. 1A), we found 4 DEGs in the JX1 samples and 0 DEGs in the 1×J samples (Table S3). We can say that the transcriptomes of all parental haploinsufficient embryos were normal, in agreement with the full potential of those embryos to develop to term and reproduce despite the initial slight developmental delay (Fig. 1H).

### Maternal EHMT2 suppresses transcriptional variation in the embryos

To reveal the long-lasting effect of maternal EHMT2 depletion in the oocyte on the transcription regulation in postimplantation development, we analyzed the RNA-seq analysis of MATMUT and control MATWT embryos at the 6-somite stage. A total of 57 DEGs were identified in this comparison (Table S3). The GO analysis of the DEGs (Fig. S3A and Table S6) revealed GO terms including erythrocyte development and homeostasis, myeloid cell homeostasis, regulation of mesoderm development, regionalization, endodermal cell differentiation, somite development. The small number of DEGs and the variability of MATMUT samples observed in the heatmaps (Fig. 5 A and B) prompted us to look for variance in the data. We calculated the average variance across genotypes and found that the variance was greater in the MATMUT embryos compared to the MATWT controls among both upregulated and downregulated DEGs (Fig. S3B). The PCA for the downregulated genes (Fig. S3C) showed greater scattering of the MATMUT samples than the MATWT samples in the first two principal components. Using MDSeq, we identified the set of genes, which were significantly (FDR < 0.05) differentially variable between the MATMUT and MATWT control embryos (Table S7). The heatmap (Fig. S3D) shows the Z-scores of the differentially variable genes in the MATMUT, and MATWT, embryos, among other samples of the current analysis. We performed a GO analysis on the set of differentially variable genes, and displayed the ten most significant terms in the bubble plot (Fig. S3E). Interestingly, the pathways again included erythrocyte homeostasis and differentiation and myeloid cell homeostasis (Table S6). Genes, dispersed in the MATMUT embryos, appeared to change developmentally between the 6-somite and 12-somite stages irrespective of mutant status. This is interesting, and tells us that the maternal mutation increases the variation at a set of genes that change during normal development. We examined some of the genes that showed variability in zygotic mutant HOMO6 embryos and found that those were also variable in the MATMUT embryos (Fig. S3F-H).

### EHMT2 regulates transposable elements predominantly in the “what it takes to be normal” category

It is known that H3K9 methylation is required for suppressing transposable elements (TEs) in the genome. Such suppression has been shown in the germ line and in ES cells (26, 27, 29, 31). We asked whether EHMT2 regulates TEs in the embryos at 6-and 12-somite stages. We identified differentially expressed (DE) repeats using uniquely mapped reads in our datasets and classified the DE repeats according to our four-way comparison (Fig. 6A and B and Table S8). We found that most DE repeats belong to the “what it takes to be normal” category. Repeats can be not only downregulated (Fig. 6A and C) but also upregulated (Fig. 6B and D) in the WT versus HOMO embryos. Specifically, 744 repeats showed downregulation in the intersection of WT12-HOMO12 and WT6-HOMO6 comparisons (orange arrow in Fig. 6A), suggesting that these were suppressed by EHMT2 in the WT condition, irrespective of somite number. These belonged to many different repeat classes (Fig. S4A and Table S8). Unexpectedly, 265 TEs were upregulated in the intersection of WT12-HOMO12 and WT6-HOMO6 comparisons (orange arrow in Fig. 6B), suggesting that these elements require EHMT2 for activation in the WT condition irrespective of somite number. These 265 repeats belonged predominantly to the LTR class (Fig. S4B and Table S8). Furthermore, 333 TEs showed downregulation in the WT12-HOMO12 but not WT6-HOMO6 comparison, and 308 repeats showed the opposite (Fig. 6A and Fig. S4C). Also, 225 TEs showed upregulation in the WT12-HOMO12 but not WT6-HOMO6 comparison, and 405 repeats showed the opposite (Fig. 6B). SINE elements were frequently found in these 405 repeats (Fig. S4D and Table S8).

**Figure 6.**
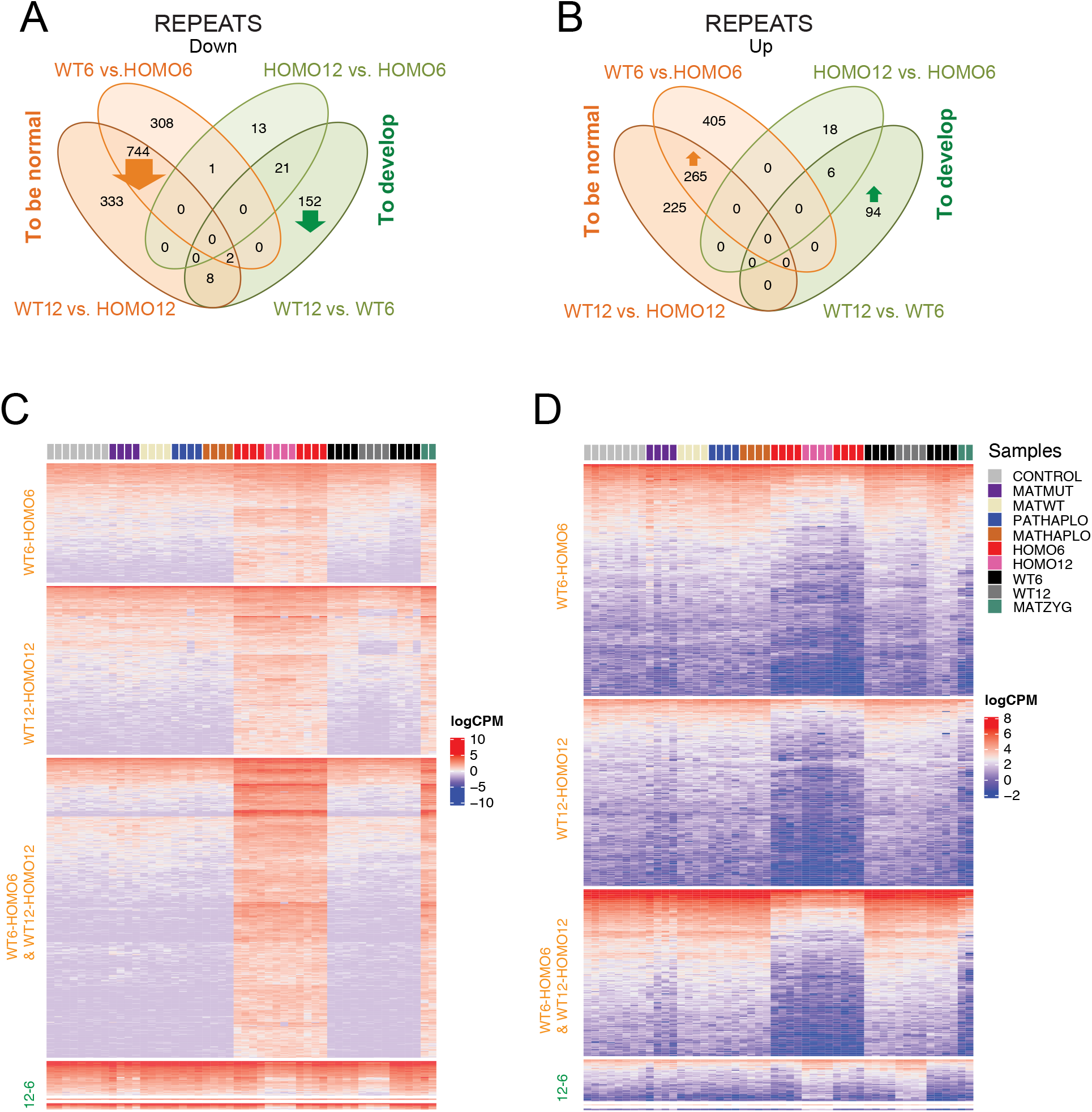
Transposable elements are misregulated in the *Ehmt2* mutant embryos. (**A-B**) Venn diagram showing downregulated (**A**) and upregulated (**B**) repeats were identified using uniquely mapped reads and were stratified according to the four-way comparison. (**C-D**) Heatmaps of downregulated (**C**) and upregulated (**D**) repeats as shown in the above Venn diagrams, with comparisons labeled to the left, samples indicated at the top and color codes shown to the right. The category labeled 12-6 contains WT12-WT6 and HOMO12-HOMO6 and their intersection.

We next determined the DE repeats that belonged to the “what it takes to develop” category (Fig. 6 and Table S8). We found that 152 repeats (mostly SINEs located at intergenic regions) were suppressed and 94 repeats (mostly LINEs) were activated in WT12 compared to the WT6 condition (Fig. S4E-F ad Table S8). The majority of the changes occurred in the “what it takes to be normal” category, suggesting that repeats depend on EHMT2 for the suppression or activation of those repeats in the embryo irrespective of developmental stage. Fewer expression changes occurred between the 6-and 12-somite stages, suggesting that TEs are not changing much during this period of embryo development, and the few changes that take place are less affected by EHMT2. It was interesting to note that the WT12-WT6 DE repeats displayed the greatest variation at the 6-somite stage, similar to WT12-WT6 DE genes. This suggests that EHMT2 regulates variation not only of genes but also of TEs.

### *Ehmt2* mutation affects distinct TE classes differently

Because some repeat classes were found more frequently in specific comparisons, we wondered whether EHMT2 has distinct roles at different TE classes. We examined the genome-wide expression of TE classes in our dataset (Table S9) and found that this is true for a few classes. As illustrated by heatmaps, the MMETn-int, IAP-d-int and MurSatRep1 TE classes were misregulated in HOMO samples irrespective of somite number (Fig. S5A-D). We detected a cluster of zinc finger protein genes that are driven by alternative promoters located at MurSatRep1 repeats and require EHMT2 for their activation in WT6 and WT12 embryos (Fig. S5D). On the other hand, some TE classes changed expression during normal development independent of EHMT2, for example, the RLTR1B-int repeats are downregulated from the 6- to 12-somite stage irrespective of genotype (Fig. S5E-F).

### EHMT2 suppresses long noncoding transcripts driven by transposable elements

When we sorted the misregulated repeats by chromosomal location, we noticed large clusters of many different kinds of repeats following the same misexpression pattern, for example of being derepressed in HOMO embryos (Table S8). We found that each cluster of misexpressed TEs initiated in one driver TE, which displayed a DMR between WT and HOMO embryos. The result of this analysis is displayed in Figure 7A. We show TE-DMRs in WT and HOMO embryos and transcription along 10 kilobases before and after each TE. We found that the ERVK TE family contributed 70% of the 145 “driver” DMR-TE-s (Fig 7B). One particular TE from the ERVK family, called RLTR17 alone made up 22% of the DMR TEs. However, not all RLTR17, but only a small fraction (18/1739) were hypomethylated in HOMO embryos. We searched for potential transcription factor binding sites in the RLTR17s hypomethylated in HOMO embryos versus all RLTR17s in the genome. We found that the mouse upstream binding protein 1 (UBP1) transcription factor site TCTCTGG was enriched (P= 8.36E-06) in those RLTR17s (Fig. 7C). UBP1, therefore, may help ERVK LTRs to gain DNA methylation in the presence of EHMT2 in WT embryos. The transcripts out of these DMR TE-s were often quite long, even hundreds of kilobases. We display examples of ERVK DMRs that initiate long transcripts in HOMO6 and HOMO12 embryos in Figure 7D-E. One example is close to 90 kb long and initiates in RLTR20C type ERVK DMR that is located in the *Platr4* TSS. Another example is an about 900 kb long transcript that initiates in an exon of *Fhit*, which appears to be an alternative promoter DMR and there is a closely located RLTR24 ERV1 upstream of it. The third example is a novel long transcript in chr1 that initiates from an RLTR17 ERVK in HOMO6 and HOMO12 samples. These three long transcripts (Fig. 7D) run through numerous passenger repeats of many kinds (Table S8). These results collectively demonstrate that EHMT2 is an important regulator of TEs in the embryo, and has high specificity to suppressing the young ERVKs in the embryo.

**Figure 7.**
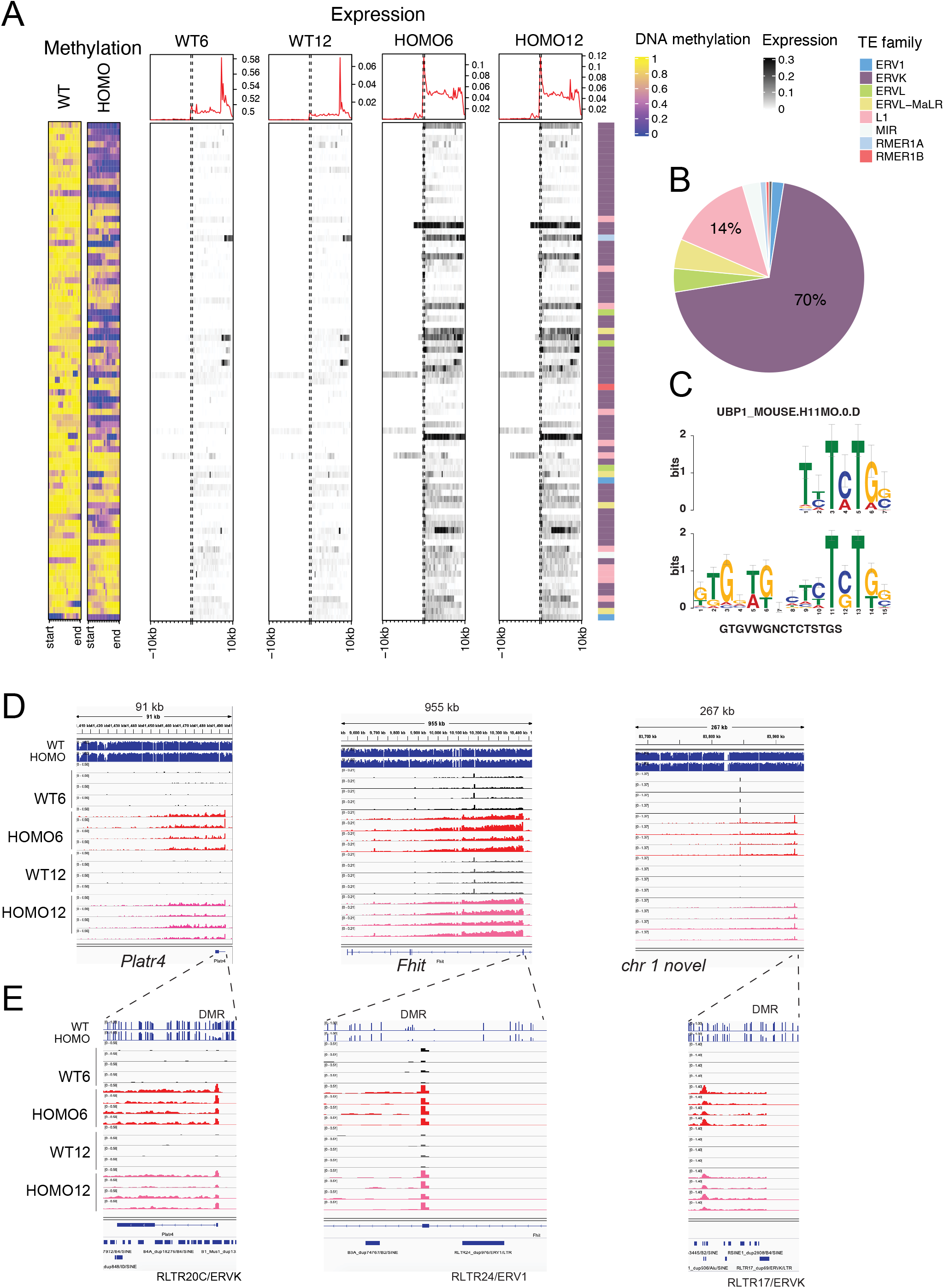
EHMT2 is required for DNA methylation targeting and suppression of long noncoding transcripts from transposable elements predominantly of the ERVK-class. (**A**) Heatmaps of differentially methylated TEs associated with differentially expressed long transcripts. DNA methylation is displayed at TE-DMRs in WT and HOMO embryos (to the left) and transcription is displayed along 10 kilobases before and after each TE in the embryos marked at the top. Normalized expression levels (logCPM) and the TE families are shown to the right. (**B**) Pie chart displays the proportion of DE repeat classes associated with DNA methylation change using the color code as above. (**C**) Known consensus sequence of the mouse UBP1 transcription factor (top) which shows enrichment (P= 8.36E-06) in RLTR17 ERVK LTRs hypomethylated in HOMO embryos (bottom) versus all RLTR17 ERVK LTRs. (**D**) Examples of ERVK DMRs that initiate long transcripts in HOMO6 and HOMO12 embryos (IGV browser images of WGBS and RNAseq experiments are displayed in the four replicate samples indicated to the left). (**E**) Magnification of the ERVKs and surrounding regions.

## DISCUSSION

The developmental progress and potential of the embryo depends on the dose of EHMT2 not only in the embryo but also in the gametes. The different severity of developmental delay in the embryos carrying different *Ehmt2* deficiencies prompted us to investigate the transcriptomes of zygotic and maternal effects and parental haploinsufficiency effects side-by-side. We asked whether delayed development correlated with changes in the transcriptome to predict the outcome of survival. We found that the transcriptome of HOMO embryos, which invariably die, is robustly distinguishable from WT embryos that reach the same somite-stage at an earlier time point. However, the transcriptomes are not altered in developmentally matched wild type MATHAPLO, PATHAPLO and WT embryos compared to CONTROL embryos. This negative result is a significant finding and suggests that these embryos develop slowly but otherwise normally after they have overcome an initial delay. Such cases likely occur during human pregnancies, when the fetus would have a slower and longer gestation despite being genetically normal. MATMUT embryos, however, that lacked EHMT2 in the oocyte, suffered lasting effects into the 6-somite stage. DEGs involved in erythrocyte and myeloid cell homeostasis were not properly suppressed and genes involved in mesoderm development, endoderm development, somitogenesis, and regionalization were not properly activated. In addition, higher variability was observed for genes in the categories of erythrocyte and myeloid cell homeostasis. We speculate that embryos, which have less variation at the 6-somite stage, have a better chance of proceeding through development. The zygotic mutation-specific pattern dominated over the maternal mutation pattern in the rare MATZYG embryos. We speculate that had they not inherited the mutant allele from the father, they would likely belong to those MATMUT embryos that are lightly affected and develop to term. Having little effect from the maternal mutation may be required for a MATZYG embryo to develop to the 6-somite stage.

We found that EHMT2 is important in reducing the variation of developmental genes after gastrulation at the 6-somite stage. One group of genes was unusual, being developmental genes, such as *Nodal, Lefty1*, Lefty2, *Cdx1*, and *Foxh1*, and showed an accelerated suppression in HOMO6 embryos. These genes play a role in breaking the symmetry, which is an essential event in the turning of mouse embryos and in development of organ asymmetries (32). Proper suppression of these genes may also be as important once their mission is accomplished, as it is to activate them at an earlier time during development. Transcriptional noise in general is an important property of normal development, and it may be important at times when major decisions are made toward cell fates. Indeed, an increased transcriptional noise was observed in the process of gastrulation by single cell RNA sequencing (33). It may be very important to curb the noise once it is no longer needed. We found that some of the developmentally changing genes displayed an increased transcriptional variation in HOMO6 embryos compared to WT6 embryos. *Nodal* and *Lefty1*, for example, exhibited high variability in their suppression at the 6-somite stage in HOMO6 embryos in contrast to their tight regulation in WT6 embryos. Our results are consistent with the role of EHMT2 in curbing the noise of developmental genes. Further experiments will address this finding at the single cell level.

The exact timing of the blockage that initiates the delay in the different *Ehmt2* deficiencies will require further experiments. The 6.25 dpc and 8.5 dpc stages are already delayed in HOMO embryos (22, 23) and the blastocyst stage may already be delayed in MATMUT embryos (2, 14). To understand transcriptional changes in mutant embryos that are delayed in development we applied two different strategies that considered the progress of embryo development. We demonstrated the usefulness of these approaches by focusing on the effects of EHMT2 in the embryo at 8.5-9.5 dpc and using somite matched embryos of different genotypes. A previous RNAseq study of *Ehmt2*^−/−^ embryos at 8.5 dpc (22) identified 253 DEGs with elevated and 181 DE genes with reduced expression in the *Ehmt2*^−/−^ mutant embryos. The authors focused on those DEGs that required EHMT2 for suppression. The large number of genes that appeared to require EHMT2 for activation in WT embryos remained unexplained in that study. Our study provides an explanation for this class of DEGs. They reflect the developmental changes from the 6- to 12-somite stage in WT embryos, highlighting the usefulness of our three-way comparison. Recently, a large set of mutant mouse lines were analyzed at 9.5 dpc by applying a similar three-way concept. Mutant embryos were contrasted against somite-staged wild type embryos from a universal baseline developmental dataset (34). This method improved the detection of real DEGs that are in direct response to the mutation. It also excluded false positive hits that corresponded to normal development. However, unlike our four-way comparison, it did not allow detecting the developmental changes that occur specifically during wild type versus mutant development, because only one time point was analyzed for each mutant line in that study. Our 4-way comparison uniquely identified developmental DEGs that require EHMT2 for their consistent suppression or activation in normal development between the 6- and 12 somite stages. The 75 DEGs that need EHMT2 for their precise suppression include genes, such as Nanog, Mesp1, Nodal, Hnf1b, that are related to gastrulation, whereas the 191 DEGs that require EHMT2 for their precise activation include genes, such as *Erbb4, Neurog2, Wnt7b, Wnt8b, Myf5*, and *Fgf9*, relevant for organ growth, connective tissue development, striated muscle development, muscle differentiation, and cartilage development. These findings establish EHMT2 as an important regulator of the transition between gastrulation and organ specification.

Our results showed DNA methylation-mediated suppression of germ cell-specific transcripts requires EHMT2 in the embryo, in agreement with an RRBS experiment (22). Our comparative analysis revealed in addition that EHMT2-dependent DNA methylation is specific to DE genes in the “what it takes to be normal” class, but not in the “what it takes to develop” class. This is consistent with the finding that developmental changes were more often up-regulations, which could not directly employ a repressor mechanism imposed by EHMT2. That same study (22) showed that DNA methylation at repeats does not globally require EHMT2 in the embryo as it does in ES cells. Similarly, maternal EHMT2 has very limited effect on DNA methylation in the oocyte and in 2-cell embryos as assessed by WGBS, but it affected two ERVKs, RLTR13D1 and RLTR46A2 (2).

Our WGBS results also found a limited effect of EHMT2 on DNA methylation at TEs globally, but we found that EHMT2 specifically targets ERVKs for DNA methylation in the embryo. ERVKs are young repeat elements with high CpG content, and are under dual control of DNA methylation and H3K9 methylation (35). We also reveal a unique feature of EHMT2-regulated ERVKs. Long noncoding transcripts could initiate in HOMO embryos from such hypomethylated repeats, and extend to several hundred kilobases, encompassing a multitude of additional, similarly misexpressed ‘passenger repeats,’ which do not display DNA hypomethylation. Transcriptional elements that retain their regulatory potential provide a rich source of functional elements in the host genome (36). We found that the hypomethylated RLTR17 ERVK LTRs were enriched in the UBP1 TF binding consensus sequence. UBP1 may recruit EHMT2 to promote DNA methylation in normal embryos. We also provided a global profiling of transposable elements specifically in the embryo. Specific repeat classes respond uniformly to EHMT2. MurSatRep1 TEs require EHMT2 for activation at a cluster of *Zfp* genes in WT embryos and RLTR1B-int TEs are downregulated during normal development from the 6-somite to 12-somite stage regardless of EHMT2. We found using the 4-way comparisons that similar to DEGs, certain TE classes correlate with genotype, while others correlate with developmental stage. Contrary to DEGs, however, differentially expressed TEs most often correlate with genotype.

Epigenetic modifiers must regulate the inevitable transcriptional noise that underlies phenotypic variation (37, 38). Maternally deposited TRIM28 is essential to carry out this role during preimplantation (39). TET1 and TET3 curbed transcriptional noise at the 8-cell and blastocyst stages (40). Our work establishes EHMT2 as a regulator of transcriptional noise after gastrulation in a developmental context. We found it interesting that curbing the noise by EHMT2 was specific to genes that switch developmental programs between gastrulation and organ specification, and also occurred at transposable elements that switched off or on between 6-and 12-somite stages. By comparing somite-stage matched individual embryos with different level of delay we were able to relate the control of transcriptional variability and the level of developmental variation. In addition, those also matched with the developmental potential of the different embryos.

### METHODS

#### Mice

All animal experiments were performed according to the National Institutes of Health Guide for the Care and Use of Laboratory animals, with Institutional Care and Use Committee-approved protocols at Van Andel Research Institute (VARI).

An *Ehmt2* conditional knockout mouse line (*Ehmt2^fl/fl^*) was generated in our laboratory by gene targeting in 129S1/SvImJ ES cells (24). The floxed SET domain was removed to generate the *Ehmt2*^-(129S1)^ allele by crossing the *Ehmt2*^*fl*/+^ male with 129S1/Sv-Hprt^tm1(CAGcre)Mann/J^ transgenic mouse female (41). *Ehmt2^fl/fl^* and *Ehmt2*^+/−(129S1)^ mice were maintained on 129S1 background. We backcrossed the *Ehmt2*^-(129S1)^ allele to the JF1/Ms (JF1) mouse strain for more than ten generations and obtained the *Ehmt2*^-(JF1.N10)^ allele, where most of the genome has been replaced by JF1 chromosomes during meiotic recombination events except the *Ehmt2* locus and its close neighbors (in the approximate interval of chr 17:34647146-35241746), as confirmed in the allele-specific expression analysis in 12-somite embryos.

*Ehmt2*^−/−^ and WT embryos were obtained by natural mating of *Ehmt2*^+/−^ females with *Ehmt2*^+/−^ males. Having *Ehmt2*^+/−^ mice in both 129S1 and JF1 background allowed us to collect samples from reciprocally crossed F1 hybrids. Crossing *Ehmt2*^+/−(JF1.N10)^ females with *Ehmt2*^+/−(129S1)^ males yielded *Ehmt2*^-(JF1.N10)/-(129S1)^ zygotic mutant and *Ehmt2*^+(JF1)/+(129S1)^ wild type embryos. The reciprocal cross resulted in *Ehmt2*^-(129S1)/-(JF1.N10)^ mutant and *Ehmt2*^+(129S1)/+(JF1)^ wild type (biparental haploinsufficient) embryos. Control *Ehmt2*^+(129S1)/+(JF1)^ embryos were obtained by crossing wild type 129S1 and JF1 male mice.

*Ehmt2*^+/−^ (JF1.N10) females were crossed with wild-type 129S1 males to generate maternal haploinsufficient wild type *Ehmt2*^+ (JF1)/+ (129S1)^ embryos. *Ehmt2*^+/−(129S1)^ males were crossed with wild-type JF1 females to generate paternal haploinsufficient wild type *Ehmt2*^+(JF1)/+ (129S1)^ embryos. Control *Ehmt2*^+(JF1)/+(129S1)^ embryos were obtained by crossing wild type JF1 female and 129S1 male mice.

*Ehmt2*^fl/fl^; *Zp3-cre^Tg/0^* females were crossed with wild type JF1 males to generate *Ehmt2*^*mat*−/+*JF1*^ maternal mutant embryos. Their control came from crossing *Ehmt2^fl/fl^* females with wild type JF1 males. *Ehmt2*^fl/fl^; *Zp3-Cre^Tg/0^* females were crossed with *Ehmt2*^+/−(JF1.N10)^ males to generate *Ehmt2^mat-/zyg-(JF1)^* maternal-zygotic mutant embryos.

Embryos were dissected at 8.5 dpc and 9.5 dpc, and were genotyped for *Ehmt2* mutation status and for sex by PCR using their allantois as described (24).

#### RNA isolation

RNA was isolated from individual embryo samples using RNA-Bee (Tel-Test) extraction followed by isopropanol precipitation. Genomic DNA was removed with the DNA-free Kit (Ambion).

#### RNA sequencing

Total RNA-seq libraries were prepared from 500 ng total RNA with the KAPA Stranded RNA-Seq Kit with RiboErase (Kapa Biosystems, MA), and were sequenced in the Illumina NextSeq 500 platform with paired-end 75 bp read length. Paired-end reads were aligned to the mm10 genome, using STAR (v 2.6.0c) with parameters --outFilterType BySJout --outFilterMultimapNmax 20 --alignSJoverhangMin 8 --alignSJDBoverhangMin 1 -- alignIntronMax 1000000 --alignMatesGapMax 1000000 --outFilterMismatchNmax 999 --twopassMode Basic --chimSegmentMin 20 --alignIntronMin 20. Using the ensembl gtf file (version 82), read counts per transcript was generated using featureCounts (-P -s 1 --primary –Q 20 –ignoreDup). Using the Ensembl gene annotation file (v82), counts per gene annotation were generated using featureCounts. Genes with at least CPM > 1 in at least 2 of the samples were retained for analysis. Differential expression was performed using edgeR (42) and differentially expressed genes were identified using the cutoff values of FDR < 0.05 and the absolute value of the logFC > 1.2. Venn diagrams were generated using http://bioinformatics.psb.ugent.be/webtools/Venn/. Enriched gene ontology terms were determined using the “clusterProfiler” package (43) in R. A term was defined as enriched applying the adj.pvalue < 0.05. Gene Ontology was performed using the “clusterProfiler” package in R. PCA plots were denerated based off the Venn Diagram DEGs. Uniquely mapped reads were extracted (samtools view -f 255), featurecounts was used to count reads per repeatmasker (44) annotation, and differential repeat analysis was performed using edgeR. A separate analysis for variation was done using the differential expression data obtained by edgeR. For each directional change, the average variance across each genotype was plotted as a bargraph using the ggplot2 package available in R. Principle component analysis was performed on DEGs using the prcomp function in R. The first two PC’s were plotted using the ggfortify package in R.

#### Differential dispersion analysis

A differential variability analysis was performed using MDSeq (30) to determine differentially variable genes within each comparison (WT12 vs WT6, HOMO12 vs HOMO6, WT6 vs HOMO6, and WT12 vs HOMO12). Differential variation was defined as a FDR.dispersion < 0.05 and a logFC > abs(1.5). Similarly, a gene was determined as differentially variable between the MATMUT and MATWT using MDSeq, and cutoff value of FDR dispersion < 0.05. The heatmap was generated using the Complexheatmap (45) package in R. GO enrichment was performed on the differentially variable genes using the clusterProfiler (43) package. The most significant terms were plotted using the ggplot2 package in R.

#### Whole genome bisulfite sequencing (WGBS) analysis

WGBS analysis was used to map DNA methylation at 9.5 dpc. One sample contains DNA from two *Ehmt2* homozygous mutant embryos (one male and one female combined), and the control sample contains DNA from a wild type female embryo. In order to generate the whole genome libraries with bisulfite converted DNA, embryo DNA was sonicated to approx. 150 bp DNA fragments. Further, DNA was end repaired by using End-It-DNA End-Repair Kit (Epicentre) and linker ligated with T4 ligase (NEB). The linked ligated DNA was bisulfite converted using EZ DNA Methylation-Gold^™^ Kit according to manufacturer’s instructions (Zymo Research) and amplified with Pfu Turbo polymerase (Agilent). Sequencing was performed in the Illumina HiSeq 2500 platform at the Integrative Genomic Core at the City of Hope Cancer Center. Pair-end reads were aligned to the mm10 genome using Biscuit and duplicates were marked using Picard’s MarkDuplicates. DNA methylation and genetic information were extracted using biscuit’s pileup and CpG beta values were extracted using vcf2bed. Significant DMR at the TSS +/− 1000 bp were determined using metilene with options –mode 2. A custom R script was used to determine genes that overlap with significant TSS +/−1000 bp locations. Differently methylated regions were identified using metilene v0.2-8 with parameters – mode 2. A region was identified as differentially methylated when it contained at least 12 CpG-s and reached the Benjamini-Hochberg adjusted p-value of *P*< 0.05. Significant regions in the mm10 genome were annotated using Homers annotatePeaks.pl.

## ACKNOWLEDGEMENTS

We thank the Van Andel Institute (VAI) Vivarium for mouse maintenance and husbandry and the VAI Genomics Core for RNA deep sequencing of the RNA samples and the Integrative Genomic Core of the City of Hope for performing the WGBS. Funding: This work was supported by VAI. We thank Xiwei Wu (City of Hope) for analysis of the WGBS samples at the early phase of this work, and our colleagues at VAI, Tim Triche for advice in bioinformatics and Gerd Pfeifer for critically reading the manuscript.

## Author contributions

Conceptualization (PES); Formal analysis (NP, WZ); Funding acquisition (PES); Investigation (TZ, NP, JL, PS, PES); Methodology (PES); Supervision (PES); Validation (TZ, NP, PES); Visualization (NP, PES); Writing-original draft (PES); Writing-review and editing (TZ, NP, JL, WZ, PS, PES).

## Data accessibility

The sequencing results have been deposited in the GEO database with accession numbers GSE156538 and GSE156539.

## Supplemental Figures

**Figure S1.**
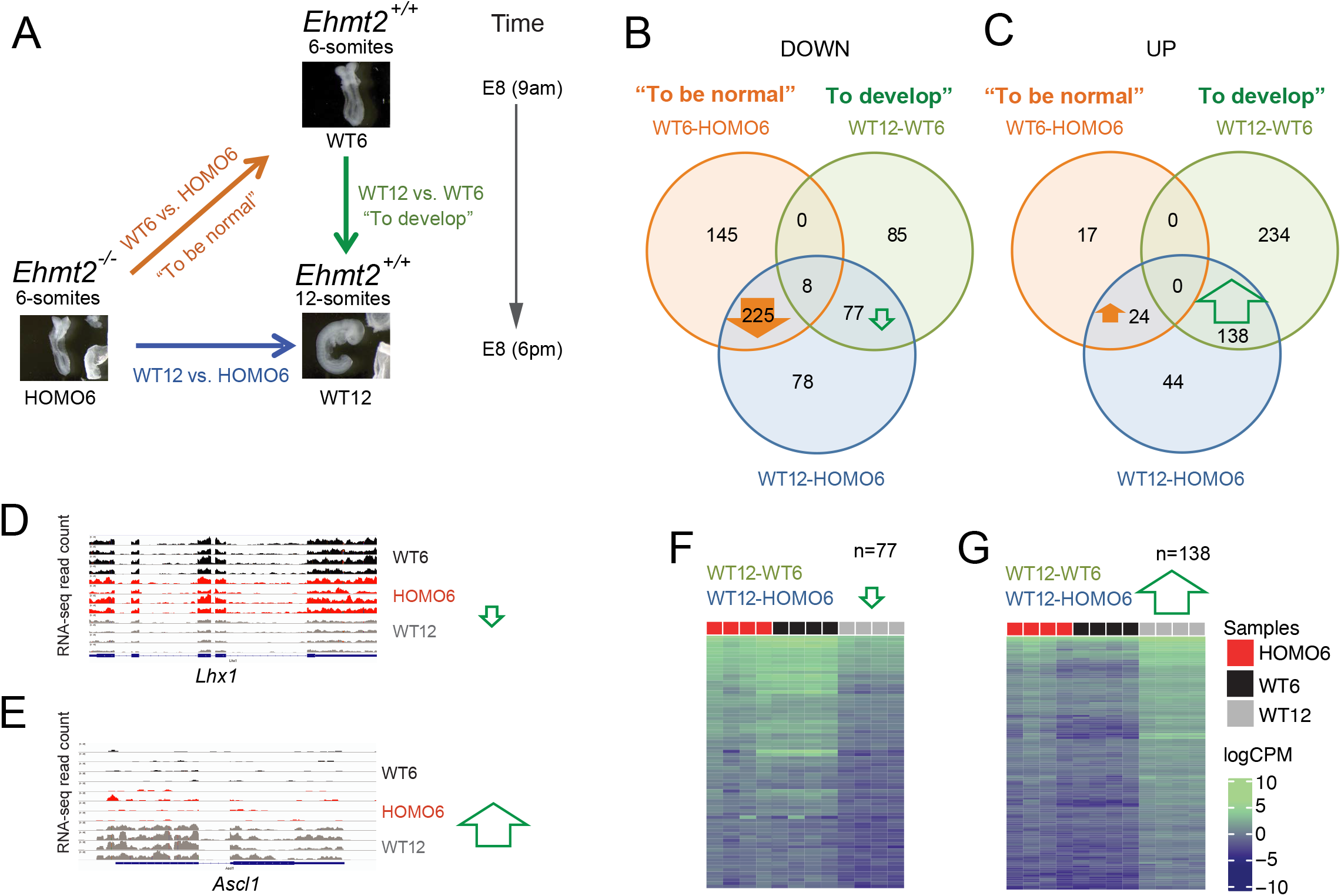
A three-way comparison reveals the direct effects of zygotic *Ehmt2* mutation and identifies false positives of the conventional method that compares siblings. (**A**) Three-way comparison. The conventional comparison (blue arrow) is normally made between uterus-mate mutant and wild type embryos. Because the mutant is developmentally delayed, the differences between these uterus-mates can be divided into two components: 1) “what it takes to be normal” (orange arrow), and 2) “what it takes to develop” from stage one to stage two (green arrow). We illustrate this idea by comparing 6-somite and 12-somite *Ehmt2*^−/−^ (HOMO6 and HOMO12) and 12-somite *Ehmt2*^+/+^ (WT12) embryos. (**B-C**) Results of the RNA-seq experiment are displayed after a 3-way comparison. Venn diagrams depict the number of DEGs along the blue, orange, and green arrows inside the circles of the matching color. (**B**) Genes are downregulated in the condition where the arrow points. EHMT2 is required to suppress 225 genes in WT6 and WT12 embryos (thick arrow). (**C**) Genes are upregulated in the condition where the arrow points. 138 DEGs require EHMT2 according to the WT12-HOMO6 comparison. Because these are also required to reach the 12-somite stage from 6-somite stage in the WT embryos (WT12-WT6 comparison), these DEGs are false positives in the conventional comparison. (**D-E**) IGV browser views are shown of normalized RNA-seq reads at false positive DEGs with reduced (*Lhx1*) or increased (*Ascl1*) expression from WT6 to WT12. (**F-G**) Heatmaps of the false positive DEGs in four samples per condition show that 77 genes are downregulated (**F**) and 138 genes are upregulated (**G**) at the 12-somite stage. Expression values (logCPM) are color coded to the right.

**Figure S2.**
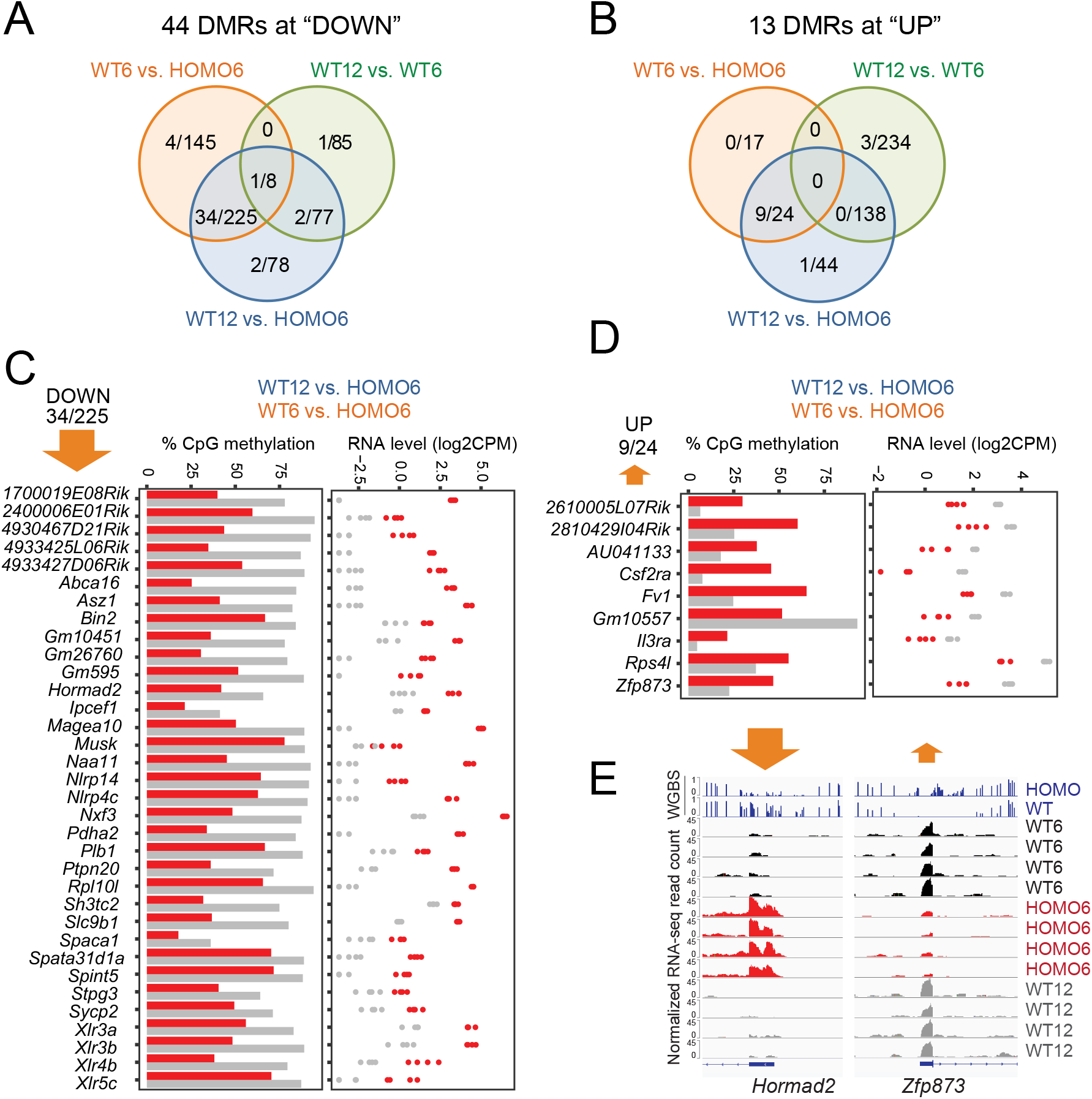
DNA methylation changes occur at a specific subset of DEGs. Venn diagrams (**A** and **B**) depict the number of DEGs, which are differentially methylated. WGBS analysis identified 57 DMRs in 9.5 dpc WT and HOMO embryos at the TSS of DEGs. (**C-D**) CpG methylation level (left) and RNA expression level (right) are plotted at DEGs with TSS DMR from the most populated sections of the Venns in **A** and **B**. Some require EHMT2 for DNA methylation at the TSS and for gene silencing (**C**), others require it for the TSS hypomethylation, and expression. Grey, WT; red, HOMO. (**D**). (**E**) IGV browser views are shown of DNA methylation and RNA expression levels at DEGs with reduced (*Hormad2*) or increased (*Zfp873*) DNA methylation in the mutant.

**Figure S3.**
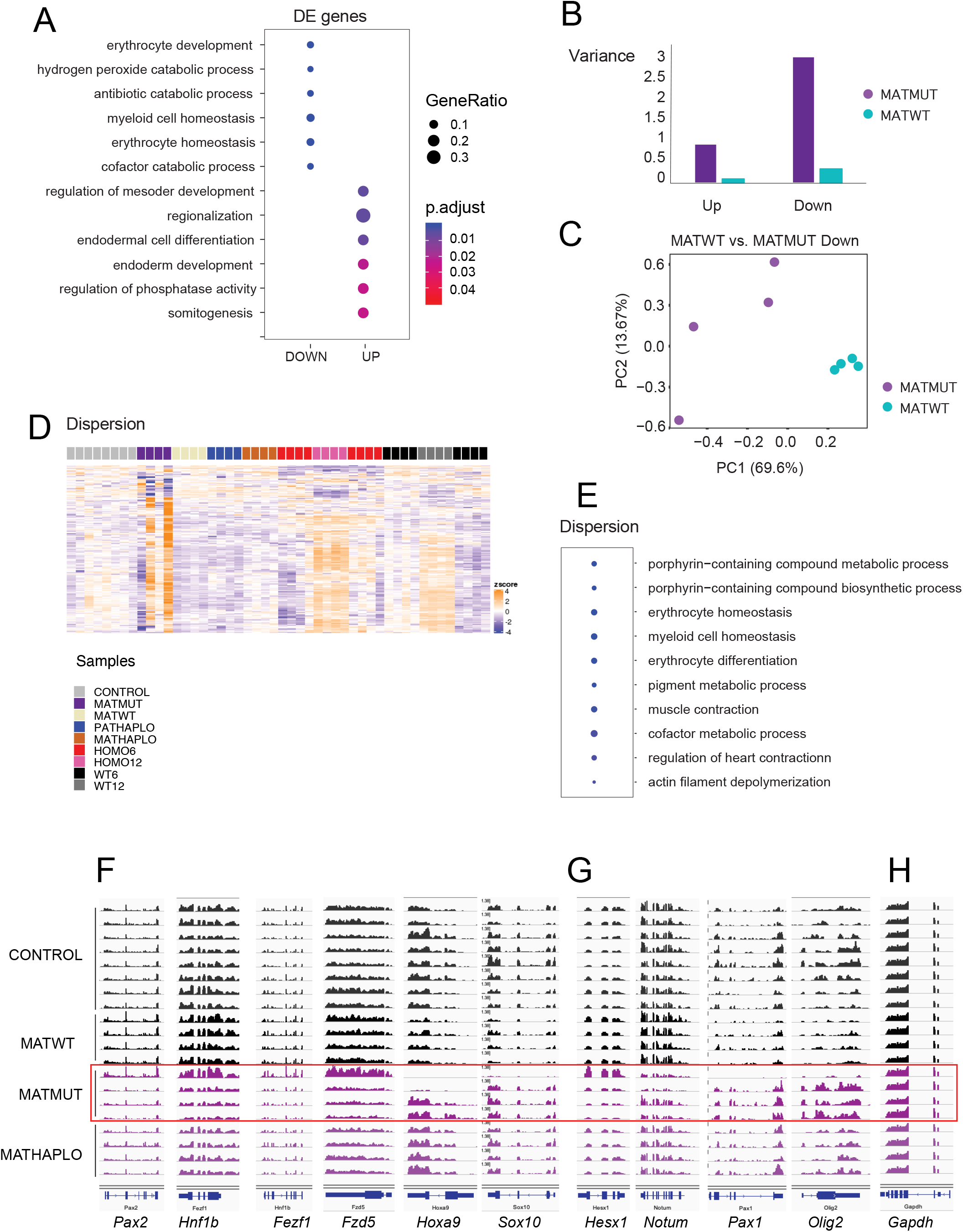
Maternal *Ehmt2* mutation results in an increased transcriptional variation. (**A**) Bubble plot from gene ontology analysis shows the most significantly enriched biological processes derived from the DEGs of MATZYG embryos compared to MATWT embryos. (**B**) Bar graph depicts the variance of DEGs identified between *Ehmt2*^*m*−/z+^ (MATMUT) embryos and control (MATWT) embryos at the 6-somite stage. (**C**) Principal component analysis of the DEGs downregulated in MATWT compared to MATMUT embryos. (**D**) Heatmap of the 500 genes that exhibit the highest level of dispersion in the MATMUT-MATWT comparison. The z-scores are displayed across the full set of samples. Note that most of these genes that are variable in MATMUT embryos show variation also between the 6- and 12-somite stages. (**E**) Gene ontology analysis depicts the biological processes derived from the most variable genes in MATMUT embryos. (**F-G**) Developmental DEGs defined in the four-way comparison also exhibit high transcriptional variation in MATMUT embryos. IGV browser images of representative DEGs are shown, which depend (F), or do not depend (G) on zygotic EHMT2 expression. (H) The control gene *Gapdh* shows no variation.

**Figure S4.**
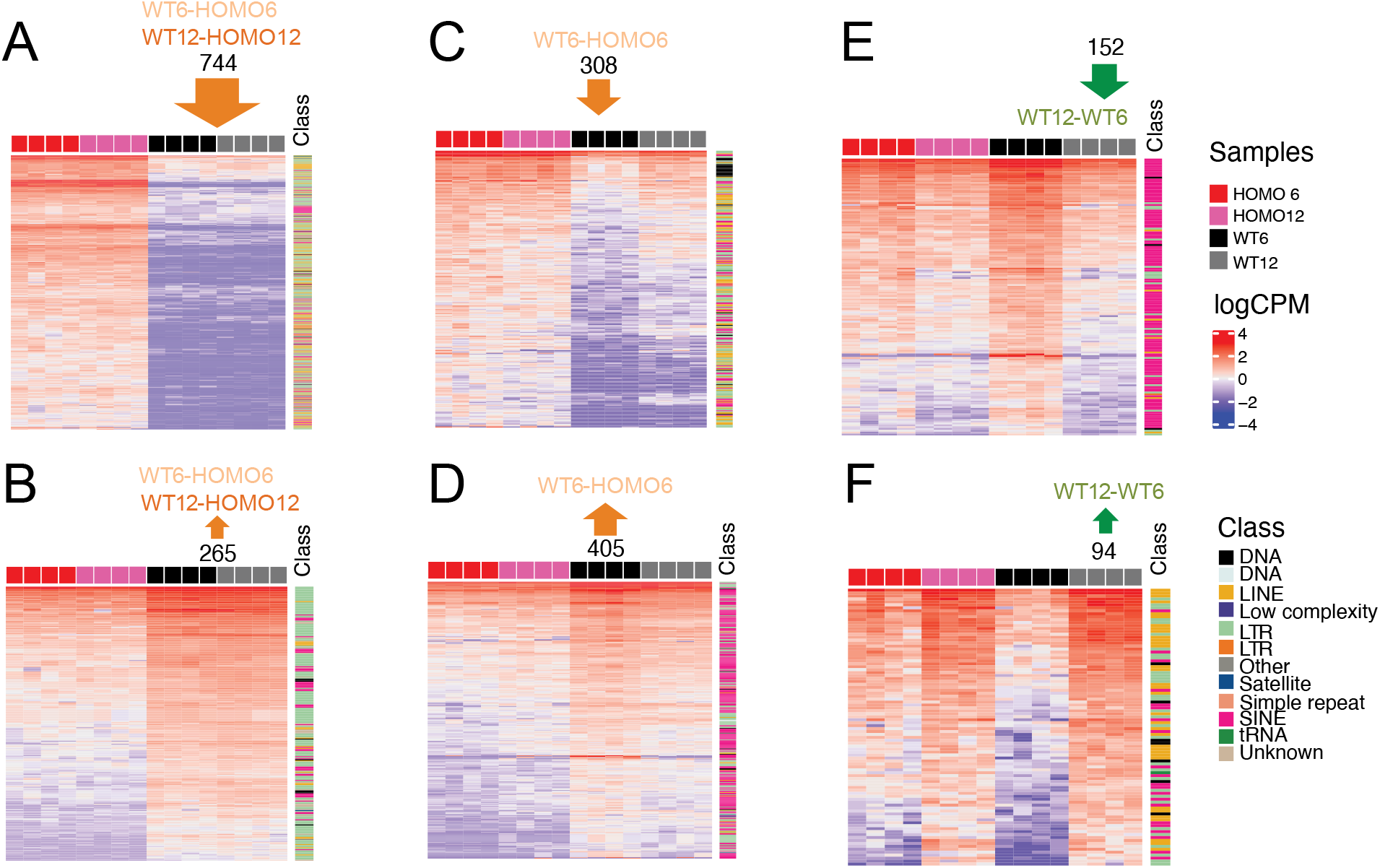
EHMT2-regulated repeat elements identified by the 4-way comparison. DE repeats were identified in the four-way comparison. Heatmaps display the transcription level (logCPM) of uniquely mapped repeats that are differentially expressed in the comparisons indicated at the top. Samples are shown by the color code to the right. The classification of each DE repeat is indicated by color code to the right. Note that variation is not suppressed in HOMO6 embryos in the WT12-WT6 comparisons.

**Figure S5.**
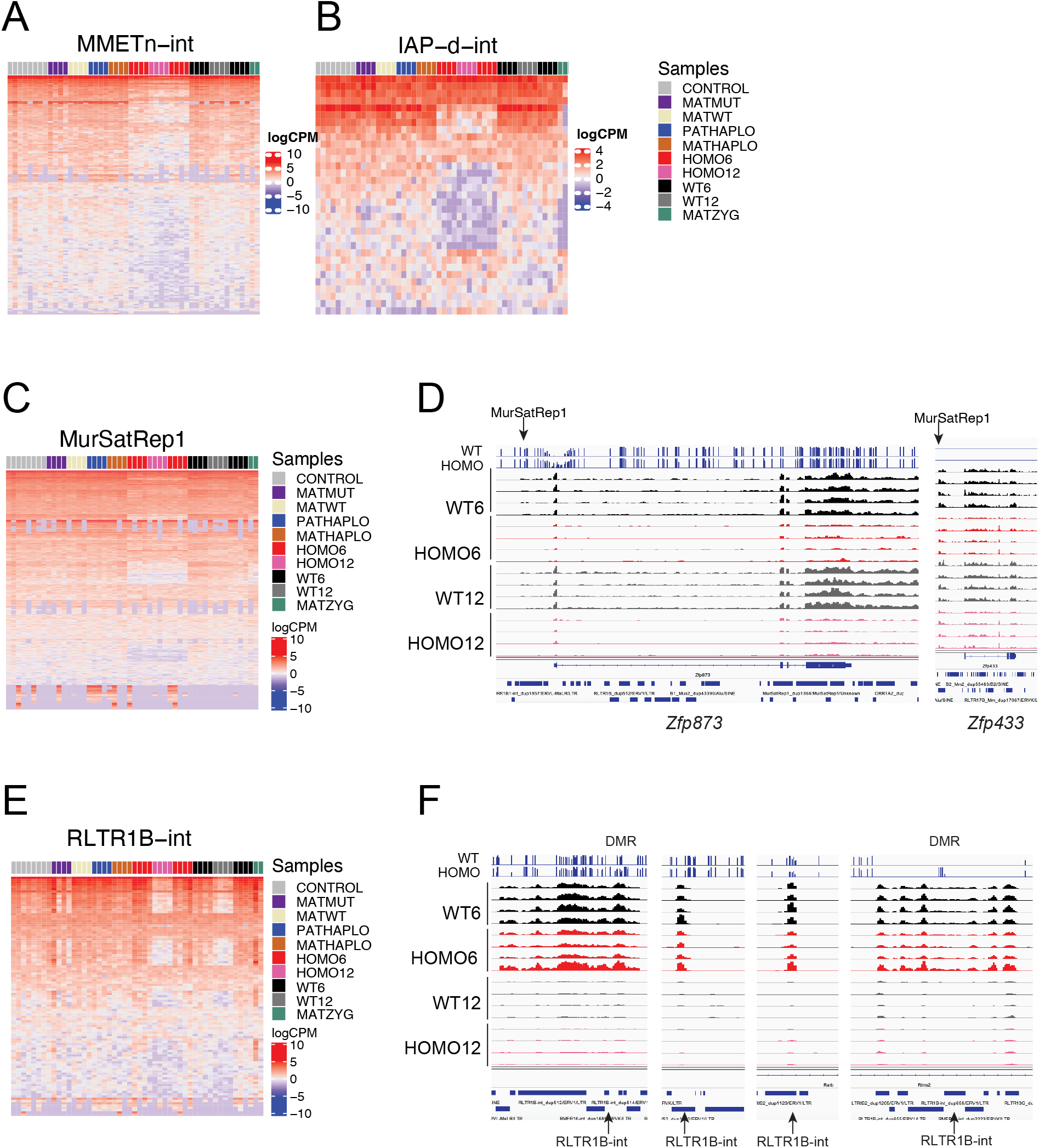
*Ehmt2* mutation affects TE classes differently. Genome-wide expression levels of selected specific repeat classes are shown by heatmaps. Note that some repeat classes correlate with genotype (**A-D**), others correlate with developmental stage (**E-F**). (**C-D**) Genome-wide expression of MurSatRep1 elements. (**C**) Heatmap displays the transcription level (logCPM) in the samples indicated by color code to the right. (**D**) Examples of zinc finger protein genes that require EHMT2 for their activation in WT embryos from an alternative promoter located at a MurSatRep1 element. (**E-F**) RLTR1B-int TEs are downregulated during normal development from the 6-somite to 12-somite stage regardless of EHMT2. Heatmap (**E**) and IGV browser views (**F**) are shown. Note in the heatmap that MATMUT samples also show variable downregulation of the same developmentally changing repeats.

## SUPPLEMENTARY TABLES

Table S1. Surviving pups with *Ehmt2* maternal mutation

Table S2. Summary of samples

Table S3. Differentially expressed genes

Table S4. Differential methylation at DEGs identified in the three-way comparison

Table S5. Gene ontology analysis of DEGs identified in the four-way comparison

Table S6. Gene ontology analysis of *Ehmt2* maternal mutant embryos by DEGs and dispersion

Table S7. Genes displaying significant dispersion in *Ehmt2* maternal mutant embryos

Table S8. The effect of *Ehmt2* mutation on uniquely mapped repeats

Table S9. Expression of uniquely mapped repeats by class

